# Adult Stem Cells and Niche Cells segregate gradually from common precursors that build the adult Drosophila ovary during pupal development

**DOI:** 10.1101/2020.06.25.171447

**Authors:** Amy Reilein, Helen V. Kogan, Rachel Misner, Karen Sophia Park, Daniel Kalderon

**Affiliations:** Department of Biological Sciences, Columbia University, New York, USA

## Abstract

Adult stem cell function relies on the prior development of appropriate numbers and spatial organization of stem cells and supportive niche cells. Drosophila Follicle Stem Cells (FSCs) present a favorable paradigm for understanding those developmental processes. About sixteen FSCs in an adult germarium produce transit-amplifying Follicle Cells (FCs) from their posterior face and quiescent Escort Cells (ECs) to their anterior. Both ECs and FCs also act as niche cells. Here we show that adult ECs, FSCs and FCs derive from common precursors intermingled with germline cells during pupal development, with progeny of a single precursor commonly including ECs and FSCs, FSCs and FCs, or all three adult cell types. Precursors posterior to germline cells become the first FCs and engulf a largely naked germline cyst projected out of the germarium to form the first egg chamber and set up a key posterior signaling center for regulating adult FSC behavior.

## Introduction

Adult stem cells generally act as a community, coordinating the activities of multiple individual cells to replenish tissues throughout life [1, 2]. The behavior of stem cells is invariably regulated by external signals from other cell types, known as niche cells [3–5]. It is therefore essential to establish adult stem cells in suitable numbers and niche environments during development to ensure appropriately regulated community activity in adults. Understanding the underlying developmental processes would facilitate identification of genetic developmental defects that confer deficiencies in adult tissue repair or cancer susceptibility, as well as the creation of functional organoids that are maintained by stem cells for therapeutic use, drug studies or mechanistic investigations [6–11].

The development of most terminally differentiated adult cell types results from a series of successive, definitive and largely irreversible decisions that are stabilized by potentially long-lasting shifts in gene expression and chromatin modification [12–17]. However, the development of adult stem cells has not been studied extensively and presents a number of special considerations. First, stem cells typically continue to proliferate in adulthood and often exhibit malleable relationships with their products, with the potential for marked changes in inter-conversion rates in either direction [18–20]. Does the general paradigm of definitive cell type specification at a fixed time in development still apply, despite this marked flexibility of the final product? Second, do adult tissues and the stem cells that maintain the tissue derive from separate or common precursors? Third, is there coordination between the development of adult stem cells and their niche cells, potentially serving to arrange appropriate proportions and juxtaposition of the two cell types?

Drosophila ovarian Follicle Stem Cells (FSCs) present a particularly interesting and experimentally favorable paradigm to investigate because the organization, behavior and regulation of these adult stem cells has been studied thoroughly, revealing many similarities to the important mammalian paradigm of gut stem cells [21–23]. FSCs also exhibit several interesting, potentially prototypical features. FSCs directly produce two different types of cell in adult ovaries, Follicle Cells (FCs) and Escort Cells (ECs) [23]. Both of these cell types also act as niche cells, producing key JAK-STAT and Wnt signals, respectively [23–27]. Paneth cells in the mammalian small intestine are also stem cell products and niche cells [28–30]. FSCs also provide a window into sustaining a tissue supported by two different types of stem cell. The continued production of egg chambers involves cooperation of somatic FSC products, ECs and FCs, with germline cells supplied by Germline Stem Cells (GSCs) [22, 27, 31]. The mechanisms of coordination among FSCs, GSCs and their products in the adult are currently not well understood but must rely on an organization that is established during development [32].

GSCs and FSCs are housed in the germarium, which is the most anterior region of each of the 30-40 ovarioles comprising the adult ovary (Fig. 1J) [22]. Somatic Terminal Filament (TF) and Cap Cells at the anterior of the germarium are stable sources of niche signals for GSCs and FSCs, including BMPs (Bone Morphogenetic Proteins), Hedgehog and Wnt proteins, with GSCs directly contacting Cap Cells [22, 31]. GSC Cystoblast daughters divide four times with incomplete cytokinesis to yield a progression of 2-, 4-, 8- and finally,16-cell germline cysts. Their maturation is accompanied by posterior movement and depends on interactions with neighboring quiescent, somatic ECs [33]. ECs also act as niche cells for adjacent or nearby GSCs and FSCs [22, 27, 34, 35]. A rounded stage 2a sixteen-cell cyst loses EC contacts as it forms a lens-shaped stage 2b cyst spanning the width of the germarium and subsequently recruits FCs (Fig. 1J). The earliest FCs can be recognized by expression of high levels of the surface adhesion protein, Fasciclin 3 (Fas3) [23, 36]. The stage 2b cyst rounds to a stage 3 cyst as surrounding FCs proliferate to form an epithelium and segregate specialized polar and stalk cells, which allow budding of the FC-enveloped cyst as an egg chamber from the posterior of the germarium (Fig. 1J) [37]. Polar Cells produce the JAK-STAT ligand, Unpaired, which patterns the subsequent development of FCs along the AP axis of developing egg chambers [37–39] and creates a gradient of signaling activity across the FSC domain, which is critical for FSC divisions and for FSC conversion to FCs [24, 26]. Egg chambers bud roughly every 12h in each ovariole of well-fed flies, progressively enlarge and mature towards egg formation and deposition, as they move posteriorly.

**Figure 1.**
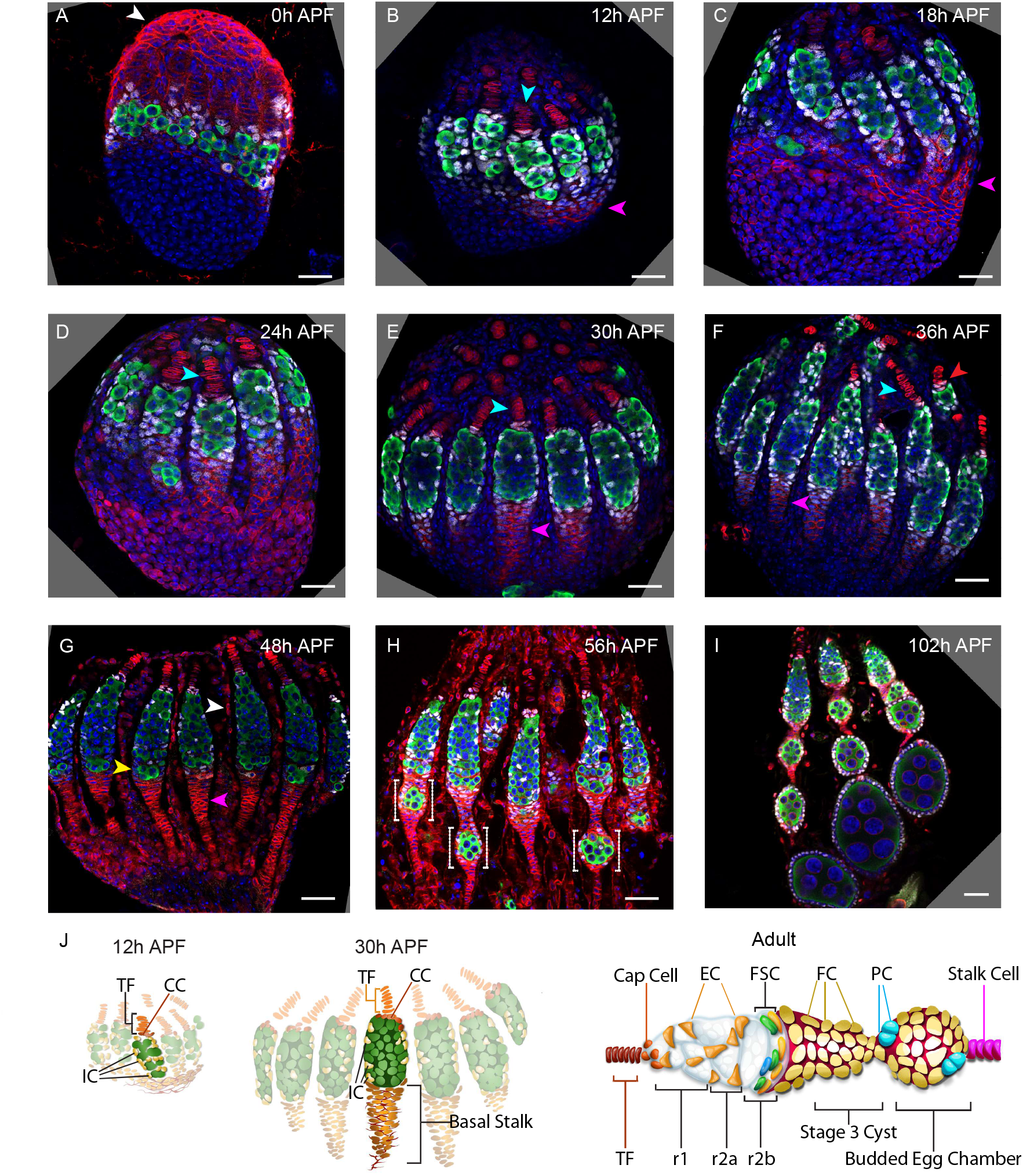
Development of adult germaria during early pupal stages. (A-J) A time-course of pupal ovary development with (J) cartoons highlighting individual germaria from 12h APF, 30h APF and adults. Vasa (green) marks germline cells, Traffic Jam (TJ, white) marks somatic ICs, DAPI (blue) marks all nuclei, LaminC (red) marks Terminal Filament (TF) cells (cyan arrowheads in D-G) and Fas3 (also red) marks basal stalk cells, except in (A) where red signal is just from Hts staining. All scale bars, 20 µm. (A) At 0h APF ovaries are not yet organized into distinct germaria. The germline (green) consists of individual cells intermingled with somatic IC precursor cells (white). The medial side of the ovary, on the right, has more developed Terminal Filaments. “Apical cells”, the precursors of epithelial sheath cells, are located at the apical (top) side of the ovary (white arrowhead). (B) At 12h APF ovaries are organized into distinct germaria, separated by a layer of migrating Apical cells. Lamin-C stains the nuclear envelope of Terminal Filament cells (cyan arrowhead). Fas3 staining is present on the medial side of the ovariole (pink arrowhead). TJ staining of somatic cells extends beyond the germline. (C) At 18h APF, basal stalks begin to form on the medial side of the ovary (pink arrowhead). (D) At 24h APF, Fas3 is spread more evenly around the circumference of the ovary, and stains a hub into which the posterior ends of the basal stalks converge. (E) By 30h APF, individual germaria are farther apart, separated from one another by a layer of 1-2 epithelial sheath cells. Lamin-C staining appears in the transition cell at the posterior end of the terminal filament, and dimly in epithelial sheath cells. (F) At 36h APF, germaria are separated by 2-3 epithelial sheath cells. Lamin-C expression is increased in epithelial sheath cells, and is also now visible (red arrow) in Cap cells (CC). Basal stalks (pink arrowheads in E-G) begin to narrow. (G) By 48h APF, the basal stalks have narrowed, and Fas3 begins to be expressed dimly in posterior ICs (yellow arrowhead). Lamin-C expression in the Cap cells increases at this stage. White arrowhead points to Lamin-C expression in epithelial sheath cells. (H) At 56h APF, germaria begin to bud the first egg chamber. (I) At 102h APF, 3-4 egg chambers have budded. (J) Cartoons illustrating individual germaria at 12h APF, 30h APF and adult stages for comparison. In the adult Escort Cells (ECs) surround developing germline cysts (white), three layers of Follicle Stem Cells (FSCs) ring the germarial circumference immediately anterior (left) to a stage 2b germline cyst and strong Fas3 (red) staining on the surface of early Follicle Cells (FCs). Polar cells (cyan) are the source of the Upd ligand that activates JAK/STAT signaling, which stimulates FSC proliferation and conversion to FCs.

There are about sixteen FSCs, which reside in three rings or layers around the inner germarial surface, between ECs and FCs on the AP axis (Fig. 1J) [22, 23]. The most posterior FSCs directly become FCs, while anterior FSCs directly become ECs, but FSCs also exchange AP locations, allowing a single FSC lineage to include both ECs and FCs [23]. Individual FSCs are frequently lost and frequently amplify, with no evident temporal or functional link between division and differentiation [40], so that the representation and output (of ECs and FCs) of any one FSC lineage is highly variable and stochastic [23]. All of these behaviors are largely regulated by extracellular signals that are graded along the AP axis, with especially prominent roles for JAK-STAT ligand derived from Polar Cells and Wnt ligands derived from both Cap Cells and ECs [23-25, 41, 42].

Previous studies have outlined how germline and somatic ovary precursors are first selected during embryogenesis, then migrate and coalesce to form a nascent gonad [43, 44]. The subsequent morphological organization of ovaries, constituent cell types and some information about relevant signals has been described up to the end of larval development [32, 45]. Understanding of pupal development is informed by a series of electron micrographs and a description of some key morphological processes [45–49] but has been limited, until recently, by technical difficulties associated with dissecting and imaging pupal ovaries [50]. Here we use morphological examination of developing pupal ovaries through fixed and live images, together with cell lineage studies to examine how adult FSCs, ECs and FCs are produced and organized from precursors present at the end of larval stages.

Marked cells derived from a single precursor clustered together along the AP axis but often spanned the border between adult EC and FSC territories. Thus, the key organizing principle guiding fate outcomes for ECs and FSCs appears simply to be a progression towards final cell locations along the AP axis without commitment to separate EC and FSC fates at a specific developmental time. The physical location and movement of precursors during the pupal period was clarified by morphological studies, including live imaging, showing that ECs and FSCs derive from precursors known as Intermingled Cells (ICs), interspersed with germline cells in the developing germarium. A more posterior group of cells gave rise to the FCs of the first-formed egg chamber. Live imaging showed that the first pupal egg chamber is formed by migration of a germline cyst out of the developing germarium into an accumulation of future FCs, rather than emergence of an FC-encased germline from the germarium that characterizes adult egg chamber budding.

The major conclusions of common precursors for FSCs and their adult niche and product cells, ECs and FCs, together with an IC origin of FSCs both directly contradict recent conclusions made from limited, targeted lineage studies [51]. We repeated those targeted lineage approaches and found results consistent with the conclusions described above. Thus, the segregation of future stem cells from niche cells in this paradigm is a gradual process, rather than a rigidly imposed demarcation of different cell types delivered at a fixed time. The juxtaposition of adult ECs and FSCs, and derivation of appropriate numbers of ECs and FSCs is assured in part by their emergence from shared precursors and from an evolving AP gradient of proliferation.

## Results

### Outline of ovary development

We dissected and imaged developing ovaries during pupation to provide a systematic descriptive timeline (Fig. 1). Germaria within each ovary develop at slightly different rates (from medial to lateral) and the overall duration of pupation can vary significantly, leading to some differences in the reported timing of critical stages [46, 52, 53]. In common with other studies, we assign “0h APF (After Puparium Formation)” as the time when larvae cease moving (pupariation) and used that event to collect animals at known intervals after pupariation. Animals mostly eclosed up to five days later (120h APF) and the first egg chamber had budded from about half of all germaria at about 56h APF (Fig. 1H).

Early ovary organization progresses from anterior (generally termed apical at early stages) to posterior (“basal”) (Fig. 1) [32, 45]. By the end of the third larval instar, each ovary has 15-20 anterior stacks of non-dividing Terminal Filament (TF) cells (Fig. 1J), which characteristically express Lamin-C but not Traffic Jam (TJ) (cyan arrowheads in Fig. 1D-F) [45, 54, 55]. “Apical cells” (white arrowheads, Fig. 1A, G), initially anterior to the TF stacks migrate towards the posterior (basally), first separating each TF stack and then entire developing ovarioles over the next 48h as they form an epithelial sheath lined by a basement membrane [46]. The most posterior “basal cells” arise from swarm cells that migrated from anterior locations prior to the late larval stage [56]. Fas3 expression initiates in a subset of basal cells on the medial side of the ovary (pink arrowheads, Fig. 1B, C) [52], spreads across the developing ovary and is then restricted to cells of epithelial appearance that intercalate progressively from posterior to anterior, to form a basal stalk (pink arrowheads, Fig. 1E-G) at the posterior end of each developing ovariole (Fig. 1J) [45, 57].

Developing germaria consist of germline cells (green in Fig. 1) and somatic cells (white in Fig.1A-I) between the TFs and basal cells. About six of these somatic cells surround the most posterior TF cell, the Transition Cell, and become non-dividing Cap Cells (Fig. 1J) [54]. Cap cells secrete Perlecan, which is essential in forming the GSC niche [58]. Cap cells also express Lamin-C and TJ shortly after the larval/pupal transition in response to Notch signaling and an ecdysone pulse [54, 59–62]. Primordial germline cells (PGCs) that contact Cap Cells largely remain in position to become adult Germline Stem Cells (GSCs) [63, 64]. The remaining PGCs develop similarly to adult Cystoblasts to produce 2-, 4-, 8- and finally, 16-cell germarial cysts during pupation.

The somatic cells posterior to Cap Cells in the developing germarium are interspersed amongst germline cells and were therefore named Intermingled Cells (ICs) (Fig. 1J). These cells express TJ (white in Fig. 1A-I) but not Lamin C or Fas3 (both red in Fig. 1B-I) at pupariation, allowing for a clear distinction from Cap Cells and basal cells [32, 52, 54]. It has generally been assumed that ICs are the source of adult ECs [32] but neither this relationship nor the source of FSCs and FCs in newly-eclosed adults has been clearly defined experimentally. Here, we use lineage analyses and the changing morphology of developing ovarioles during pupation to investigate the origin of adult ovarian somatic stem and niche cells.

### Spatially organized progression of germline differentiation

We are principally interested in understanding the developmental origins of somatic ECs, FSCs and FCs but this process must also be coordinated with germline development to produce egg chambers, starting midway through pupation. Germline development was monitored by using Hts antibody (red in Fig. 2) to reveal a rounded spectrosome, an internal organelle characteristic of single germline cells, or a branched spectrosome derivative known as a fusome, which connects Vasa-marked germline cells (green in Fig. 2) in a cyst [65], and by incubating samples with EdU to assess DNA replication as a proxy for cell division (Fig. S1).

**Figure 2.**
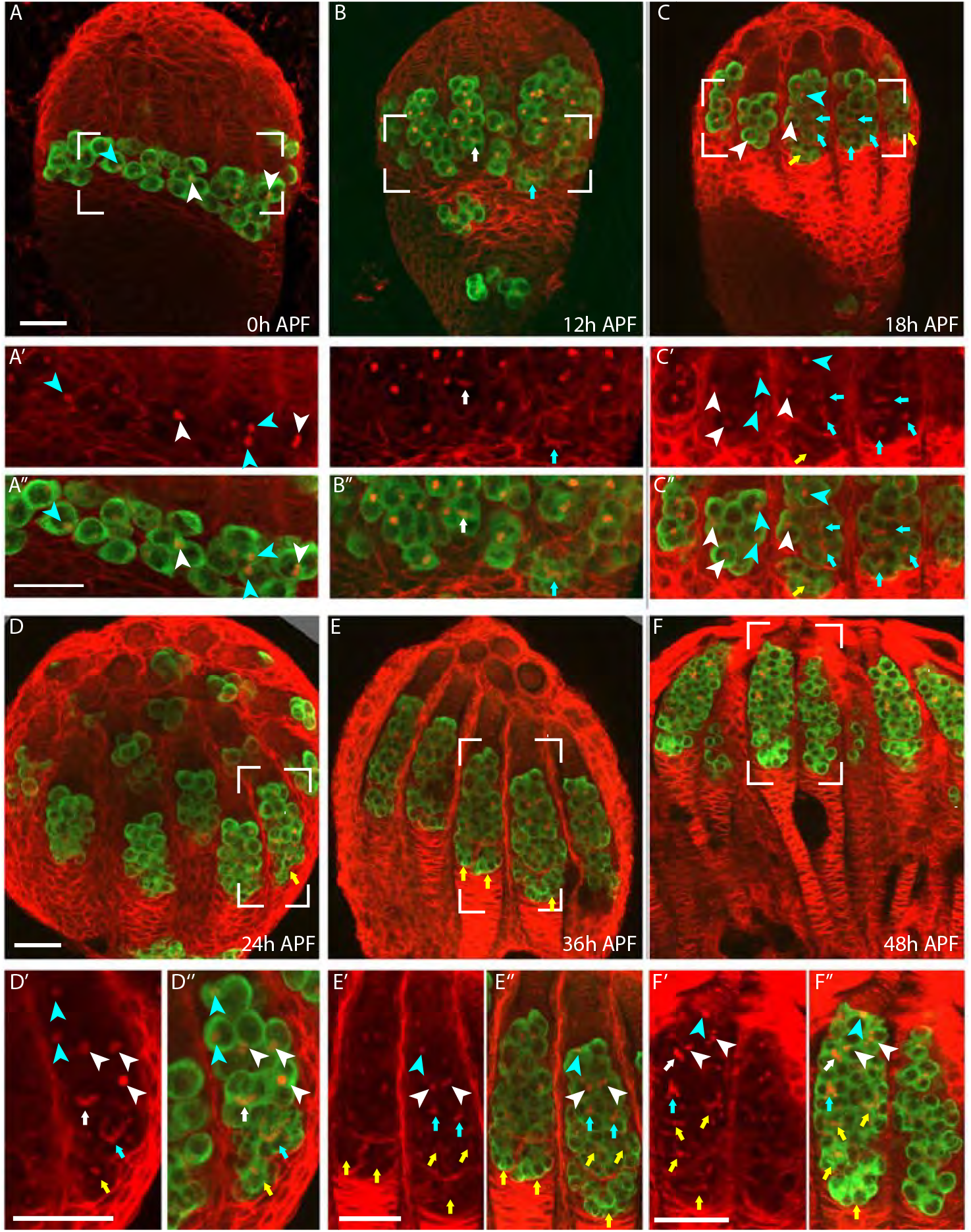
Germline development during early pupal stages. (A-F) Germline development at the indicated times after puparium formation (APF) was monitored with Hts (red), which stains spherical spectrosomes and branched fusomes in germline cells as well as the surface of many somatic cells, and Vasa (green), which identifies germline cells. The number of germline cells within a cyst was determined by examining the fusome connecting clustered Vasa-positive cells over a range of z-sections (here, single or small numbers of superimposed z-sections are shown). For each sample, a close-up of the region outlined with white brackets is shown (A’-F’) without or (A”-F”) with the Vasa fluorescence channel. (A-F) Germaria progressively lengthen from anterior (apical, top, future anterior) to posterior (basal, bottom, future posterior) as germline cells become more numerous. The number of germline cells in a cyst is indicated as one (cyan arrowhead), two (white arrowhead), four (white arrow), eight (cyan arrow) or sixteen (yellow arrow). (B, D) Some germline cells migrate through the basal cells to become stranded at the posterior of the ovary. (A-A’’) At 0h APF germline cells are single cells (cyan arrowheads) or 2-cell cysts (white arrowheads). (B-B’’) At 12h APF the majority of germline cells have 1 or 2 cells, but posterior cysts may have 4 (white arrow) or 8 cells (cyan arrow). (C-C’’) At 18h APF, there is a large range of germarial length and corresponding posterior cyst maturity. Longer germaria have 16 cells in posterior cysts (yellow arrows) whereas short germaria have only 2 cells in posterior cysts (white arrowhead). (D-D’’) At 24h APF the majority of germaria have 16 cells in posterior cysts (yellow arrows) but the most posterior cyst has not yet moved beyond other cysts. (E-E’’) At 36h APF a single cyst resides at the posterior of most germaria but in some cases the most posterior 16-cell cyst has not yet been resolved. (F-F’’) By 48h APF each germarium has a most posterior 16-cell cyst. Scale bars, 20 micron.

Primordial Germ Cells (PGCs) do not initiate differentiation before late larval stages due to the repressive actions of ligand-free ecdysone receptor and Broad transcription factor targets in somatic cells [60]. Thereafter, anterior PGCs receive BMP signals from newly-formed Cap cells that prevent differentiation, allowing them to become adult GSCs, while posterior PGCs start to express the early differentiation marker, Bam-GFP in late third instar larvae just prior to pupariation, co-incident with a strong ecdysone pulse [60, 64]. A short time later, some branched fusomes are seen in posterior PGCs, indicating divisions with incomplete cytokinesis to form cysts with 2 or more cells [53, 60]. The subsequent progression of germline differentiation during pupation has not been described.

We observed 2-cell cysts at 0h APF, cysts of up to 8 cells by 12h APF, and 16-cell cysts at 18h APF (Fig. 2A-C). Progressively larger cysts were arranged, with few exceptions, from anterior to posterior from 18h APF onwards; this was particularly clear in samples from 24h to 48h APF (Fig. 2D-F). Individual germaria within a developing ovary exhibited significantly different degrees of progression, together with AP elongation of the germarium to accommodate increasingly large germ cell units; in some cases, germline cells had also migrated posteriorly out of the germarial region, likely as a non-productive developmental aberration (Fig. 2A-D). Two or more 16-cell cysts were seen in individual germaria by 36h APF. A pair of posterior cysts were often side by side in germaria at 24h APF but only a single cyst occupied the most posterior position in most cases by 36h APF, and in all germaria by 48h APF (Fig. 2D-F). Consistent with these observations of cyst progression and location, EdU labeling was often seen in the most posterior cyst(s) up to 24h but rarely at 36h, while at 48h even the second most posterior cyst in the germarium rarely incorporated EdU (Fig. S1A-D). Our observations reveal a progression of germline differentiation that is spatially organized along the AP axis during the first 48h APF, at which time the organization of germline cysts resembles that in the adult up to a stage 2b cyst (Fig. 1J).

### Follicle Stem Cells (FSCs) and Escort Cells (ECs) derive from a shared precursor

We conducted a series of lineage analyses to learn about the specification of ECs, FSCs and FCs from precursors during pupal development. In these studies, the timing of clone initiation was deduced from the time elapsed between administration of a heat-shock to induce *hs-flp*-mediated genetic recombination and adult eclosion. Thus, a “-5d sample” was generated by collecting very young adults (less than a 6h range) centered around 120h after heat-shock (Fig. 3A). The time between puparium formation and eclosion was generally a few hours more than observed for animals used in morphological studies, but there is also a short delay between heat-shock and genetic marking of a cell [66]. Consequently, the time a cell is genetically marked in a “-5d sample” likely corresponds closely to puparium formation (0h APF) (Fig. 3A).

**Figure 3.**
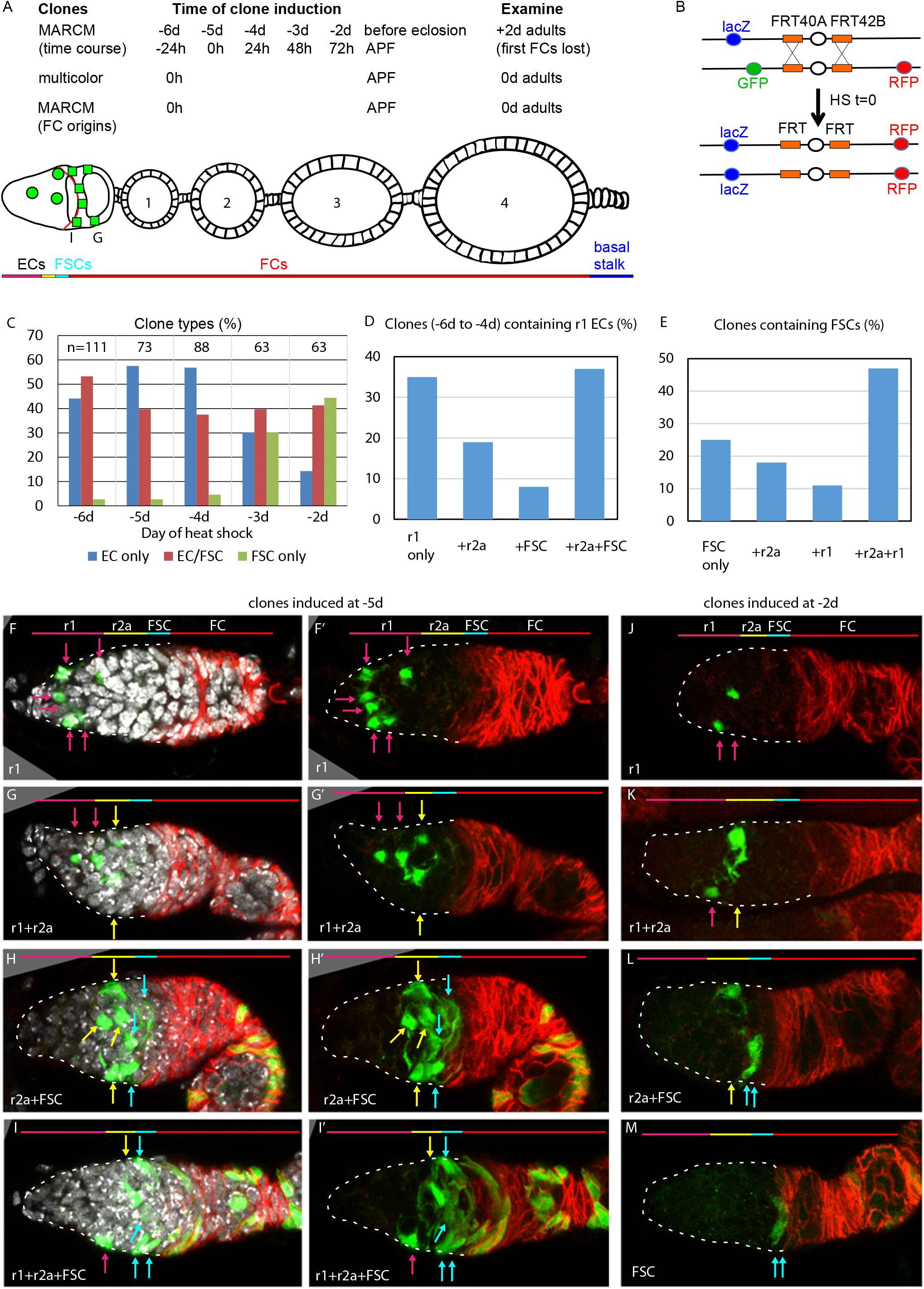
Common precursors for FSCs and ECs with derivatives clustered along the AP axis. (A) Summary of lineage tracing experiments performed in this study. The ovariole of a newly eclosed adult, which has 4 egg chambers, is depicted with an example lineage (green cells). In the first set of experiments, we dissected ovarioles from adults 2d after eclosion (“+2d”), having induced clones daily (in separate experiments) between 6d and 2d before eclosion (“-6d” to “-2d”). In these experiments the first egg chambers to bud from the germarium will no longer be present and therefore the first FCs to be made cannot be seen. Some cells that had been FSCs at eclosion will have moved into the FC region posterior to the Fas3 border (red line). To capture all labeled FCs produced during pupation we examined newly-eclosed adults. To examine the maximal proportion of marked ovarioles with a lineage derived from a single cell we used multicolor labeling. (B) Second chromosome genotype of multicolor flies used for multicolor lineage tracing of precursors, showing tub-lacZ (lacZ), ubi-GFP (GFP) and His2Av-RFP (RFP) transgenes and FRT 40A and FRT 42B recombination targets either side of the centromere (white oval). Heat-shock induction of a hs-flp transgene on the X-chromosome can induce recombination independently at either pair of FRTs, making one or both chromosome arms homozygous in daughter cells, thereby eliminating one or more of the marker genes. A possible outcome for one daughter cell is shown. The other daughter has a complementary genotype (in this case with GFP-only), so that it is generally possible to identify sister “twin-spot” lineages. (A-K) GFP-labeled MARCM clones were induced from 6d to 2d before eclosion and flies were dissected 2d after eclosion. In each germarium labeled cells were scored as region 1 (r1) ECs, region 2a (r2a) ECs, or FSCs. (A) Most clones initiated 4-6d before eclosion contained only ECs (blue) or FSCs together with ECs (red), indicating a common precursor. Clones containing only ECs declined and clones containing FSCs but no ECs (green) increased in frequency when initiated at later times. (B, C) Data were pooled from clones induced from −4 to −6d. (B) Clones containing r1 ECs (n=174) were grouped according to the additional cell types they contained; inclusion of other r1 ECs was the most frequent, and r2a ECs were more commonly included than more distant FSCs even though the total frequency of clones with r2a ECs (54%) and FSCs (57%) were similar. (C) Clones containing FSCs (n=155) contained r2a ECs more frequently than r1 ECs even though the total frequency of clones with r2a ECs (54%) was lower than for r1 ECs (64%). (D-G) Examples of different clones induced 5d before eclosion all demonstrate the clustering of marked cells (green) along the anterior (left) to posterior (right) axis. The anterior Fas3 (red) border, viewed in each z-section, allows distinction between FCs and FSCs (in the three layers anterior to Fas3); several z-sections were combined to show all labeled cells in one image. DAPI (white) staining reveals all nuclei but is omitted in (D’-G’) and (H-K) for clarity. GFP-marked clones have (D) a cluster of r1 ECs (magenta arrows), (E) a cluster of r1 and r2a ECs (yellow arrows), (F) a cluster of r2a ECs and FSCs (blue arrows), (G) r1 and 2a ECs with FSCs. (H-K) Clones induced 2d before eclosion also showed clustering of progeny, with (H) r1 ECs only, (I) r1 and r2a ECS, (J) r2a ECs and FSCs, and (K) FSCs only. Scale bars, 20 µm.

Ideally, a single precursor per developing ovariole would be labeled at a variety of times during pupation, followed by comprehensive analysis of each resultant lineage in large numbers of ovarioles from newly-eclosed adults. In practice, it is difficult to label only one cell in a developing ovariole when there are many precursors present. We used a very mild heat-shock protocol to try to accomplish this objective. We initially used MARCM clonal analysis [67] because positive cell marking with GFP greatly facilitates scoring all cells in a clone. However, maturation of a strong GFP MARCM signal in adult and developing ovarioles requires almost 4d. We therefore dissected ovarioles from adults 2d after eclosion (“+2d”) in our first experiments, having induced clones daily (in separate experiments) between 6d and 2d before eclosion (“-6d” to “-2d”). These experiments provided an important overview of the temporal progress of precursor types over the course of pupation (Fig. 3A). Subsequent experiments focused further on studying single cell lineages by using multicolor labeling to reduce the frequency of labeling a precursor in a specific color, and on capturing all labeled FCs produced during pupation by examining lineages in newly-eclosed adults (Fig. 3A)

In the first experiments, involving staged clone induction, each ovariole in 2d-old adults was scored for the number of marked FSCs (Fig. 1J), the number of marked ECs and the presence of marked FCs in the germarium and first two egg chambers (implying the presence of a FSC 2d earlier) by imaging and archiving complete confocal z-stacks (Fig. 3). Marked TF and Cap Cells were also noted. Clones were classified according to inferred cell types present at the time of eclosion (see Methods for extrapolating data from +2d adults to 0d adults).

The average number of labeled ECs and FSCs in a marked adult germarium decreased when clones were induced at progressively later stages (from an average of 8.1 at −6d to 2.0 at −2d), consistent with proliferation of most precursors during this period (Tables 1 and 2). Very few marked clones were observed in the absence of heat-shock, consistent with almost all clones arising at the time of deliberately administered heat-shock. As expected, TF cells were almost never labeled and Cap Cell labeling was seen only at the earliest times of induction (present in 3% of clones induced at −5d (0h APF)) because these non-dividing cell types have already differentiated shortly before (TFs) or just after (CCs) the larval/pupal transition and the MARCM method only labels dividing cells.

**Table 1.**
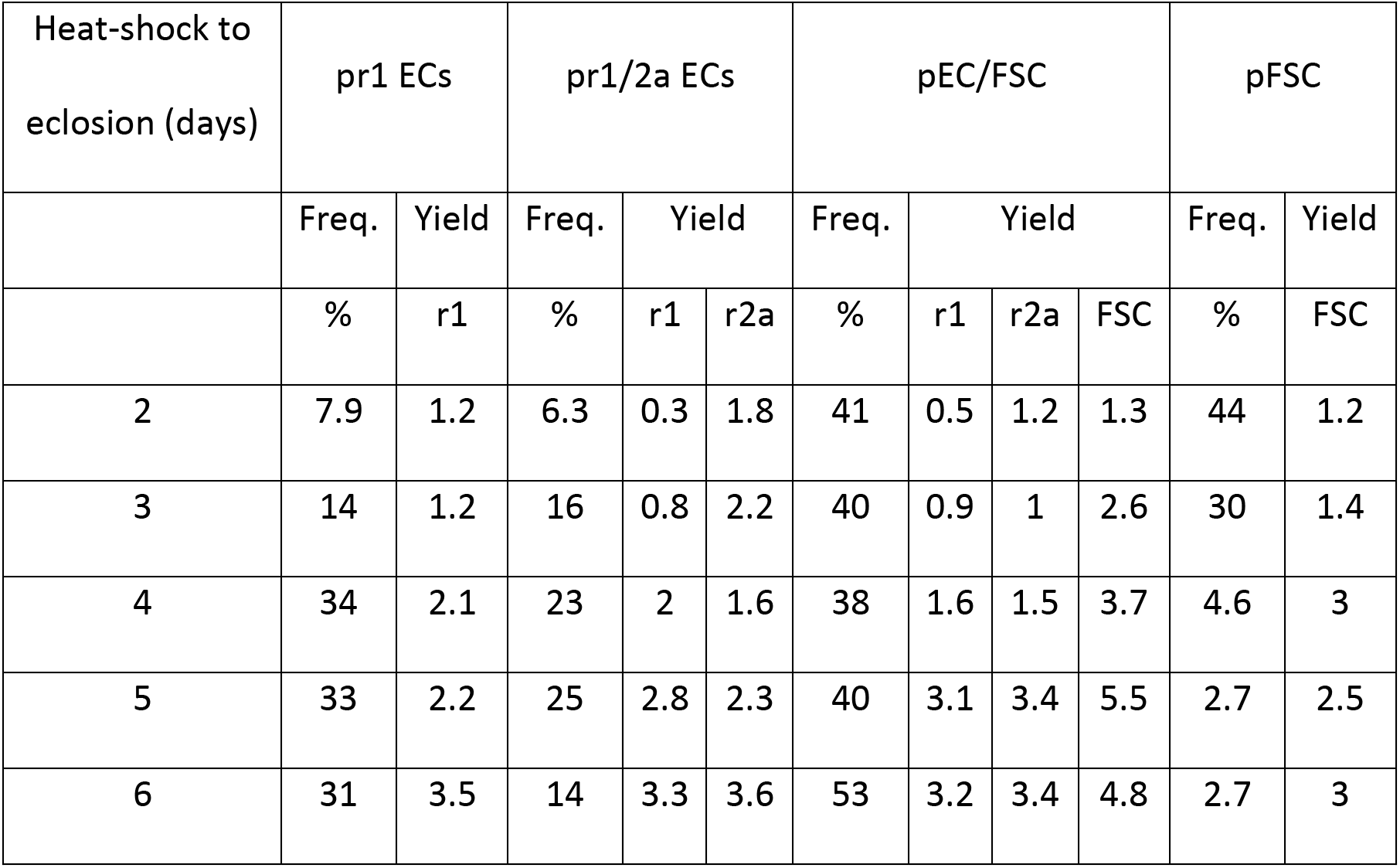
Frequency of clones originating from different precursors and yields per clone.

| Heat-shock to<br>eclosion (days) | pr1 ECs |  | pr1/2a ECs |  |  | pEC/FSC |  |  |  | pFSC |  |
| --- | --- | --- | --- | --- | --- | --- | --- | --- | --- | --- | --- |
|  | Freq. | Yield | Freq. | Yield |  | Freq. | Yield |  |  | Freq. | Yield |
|  | % | r1 | % | r1 | r2a | % | r1 | r2a | FSC | % | FSC |
| 2 | 7.9 | 1.2 | 6.3 | 0.3 | 1.8 | 41 | 0.5 | 1.2 | 1.3 | 44 | 1.2 |
| 3 | 14 | 1.2 | 16 | 0.8 | 2.2 | 40 | 0.9 | 1 | 2.6 | 30 | 1.4 |
| 4 | 34 | 2.1 | 23 | 2 | 1.6 | 38 | 1.6 | 1.5 | 3.7 | 4.6 | 3 |
| 5 | 33 | 2.2 | 25 | 2.8 | 2.3 | 40 | 3.1 | 3.4 | 5.5 | 2.7 | 2.5 |
| 6 | 31 | 3.5 | 14 | 3.3 | 3.6 | 53 | 3.2 | 3.4 | 4.8 | 2.7 | 3 |

**Table 2.** Number of adult cells of each type made from dividing precursors at different developmental times calculated from the total yield of marked cells of each type.

| Heat-shock to<br>eclosion (days) | Total Number of marked cells in<br>clones |  |  |  | Cells made per germarium from dividing<br>precursors |  |  |  |
| --- | --- | --- | --- | --- | --- | --- | --- | --- |
|  | r1 | r2a | EC | FSC | r1 | r2a | EC | FSC |
| | x | y | x+y | z | $16x/z$ | $16y/z$ | $16(x+y)/z$ | |
| 2 | 20 | 37 | 57 | 69 | 4.6 | 8.6 | 13.2 | 16 |
| 3 | 41 | 46 | 87 | 93 | 7.1 | 7.9 | 15 | 16 |
| 4 | 156 | 80 | 236 | 134 | 18.6 | 9.6 | 28.2 | 16 |
| 5 | 191 | 140 | 331 | 164 | 18.6 | 13.7 | 32.3 | 16 |
| 6 | 356 | 253 | 609 | 295 | 19.3 | 13.7 | 33 | 16 |

Strikingly, almost all ovarioles with early-induced clones (from −6d to −4d) that included marked FSCs also contained marked ECs (“EC/FSC clones”) (Fig. 3A, D-G) (Table 1). Ovarioles containing only marked ECs (“EC only”) were seen at each time of clone induction, while a high frequency of ovarioles with marked FSCs but no marked ECs (“FSC only”) was evident only when induced at −3d or - 2d (Fig. 3A, K). These observations immediately suggested that most adult FSCs derive from a dividing precursor that gives rise to both ECs and FSCs. During the second half of pupation an increasing proportion of the products of early EC/FSC precursors produce FSCs but no ECs.

### Relative location of precursors along AP axis instructs fate

We found that marked cells in an adult germarium were generally clustered in the AP dimension. This can be seen for each time of clone induction by examining individual images (Fig. 3D-K). To evaluate clustering numerically we counted marked region 1 (r1) and region 2a (r2a) ECs separately. Region 1 contains germline cysts with fewer than 16 cells and comprises the most anterior two-thirds of EC territory along the AP axis. Of all early-induced clones (−6d to −4d) with two or more marked cells and including at least one r1 EC, 35% contained only r1 ECs, 19% included also only r2a ECs, while only 8% also included only FSCs (the remainder included both r2a ECs and FSCs) (Fig. 3B). Thus, genetic relatives of a marked region 1 EC are often only in region 1; they are also more commonly found in region 2a than in the more distant FSC region, even though the total frequency of clones with marked region 2a ECs (54%) and FSCs (57%) were similar (Fig. 3B). Conversely, of all clones with two or more marked cells including at least one FSC, 25% contained no ECs, 18% included also only r2a ECs, while only 11% also included only r1 ECs, even though the total frequency of clones with r1 ECs (68%) was higher than for r2a ECs (54%) (Fig. 3C). These observations support the hypotheses that precursors occupy significantly different AP positions early in pupal development, their progeny undergo stochastic movement in either direction but still remain relatively close in the AP dimension, and cell identity (r1 EC, r2a EC or FSC) is set by their AP location at eclosion. By adulthood, this identity includes starkly different behaviors of quiescence (ECs) or proliferation (FSCs). Had precursors been restricted early in pupation to produce only FSCs, only ECs, or only one type of EC (r1 or r2a), marked lineages would be restricted to a single cell type. In fact, we observed many lineages that spanned boundaries between r1 and r2a ECs, and between ECs and FSCs, especially for clones induced from 0h to 48h APF.

### Low frequency clones validate a common precursor for ECs and FSCs

The overall clone frequency (percentage of ovarioles with any marked cells in the germarium or early egg chambers) in the above experiments was 60%. It would therefore be expected that about 2/3 of marked ovarioles derive from marking a single precursor if all recombination events were independent and equally likely (see Methods). To produce a higher frequency of ovarioles with marked lineages derived from only a single cell we used a multicolor marking strategy. Here, recombination events can lead to loss of GFP or a *lacZ* transgene, with or without RFP loss to yield six possible lineage phenotypes, markedly reducing the proportion of precursors labeled by a specific color combination (Fig3B) [23]. Two types of clone result from recombination at both *FRT40A* and *FRT42B* (GFP-only and *lacZ*-only) and are therefore the least frequent. GFP-only clones, induced at −5d and scored in newly-eclosed (0d) adults, were found at an overall frequency of 26/115 ovarioles, so that an estimated 86% of labeled ovarioles (see Methods) would be expected to have lineages derived from a single GFP-only cell.

The patterns of GFP-only and other multicolor clones were very similar to those observed by MARCM analysis (Fig. 4), with a single lineage frequently spanning r1 and r2a territory, or EC and FSC domains (Fig. 4A-C). Among GFP-only lineages that included FSCs, eight also included ECs and only two did not (Fig. 4B, C). We can therefore be confident that a single precursor at −5d (0h APF) can give rise to both ECs and FSCs and that most FSCs originate from such precursors. The proportions (and yields) of GFP-only clone types were 42% EC-only (average 3.0 ECs), 31% EC/FSC (average 3.5 EC and 2.6 FSCs) and 8% FSC-only (average 4.0 FSCs) (Table 3).

**Figure 4.**
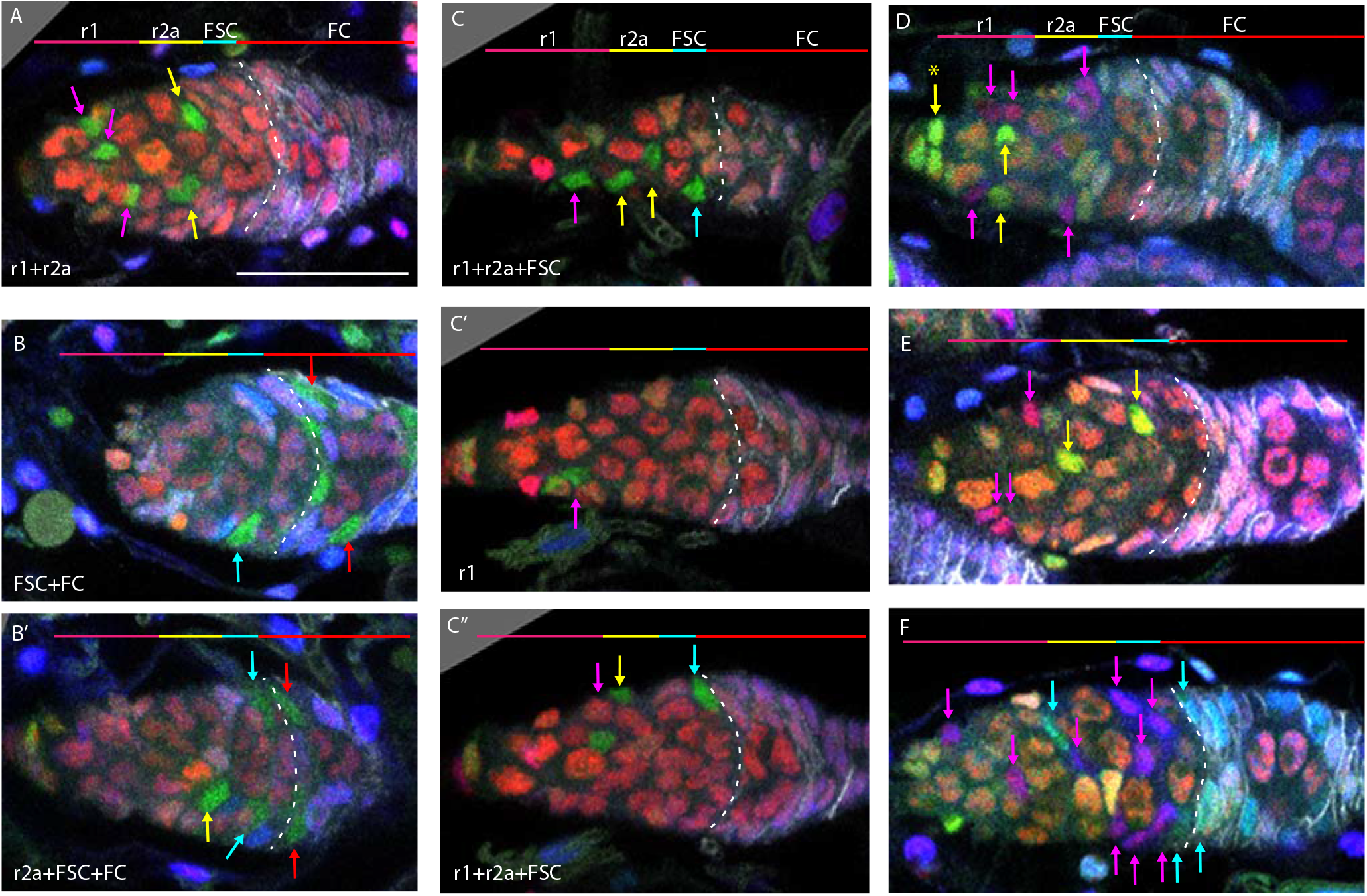
Multicolor labeling of precursors shows clustering of progeny along the AP axis. (A-F) Multicolor lineage tracing was performed by inducing clones 5d before eclosion and dissecting newly-eclosed flies. The anterior limit of Fas3 staining is marked by a white dashed line. Scale bar in (A) applies to all images, 20 µm. Precursors produce r1 (magenta) and r2a (yellow) ECs, FSCs (cyan), and FCs (red) in overlapping zones, indicated by the colored zones and in (A-C) corresponding arrow colors. (A-C) show examples of GFP-only clones (green), which are generated by two recombination events and are therefore infrequent. (A) A clone containing r1 and r2a ECs. (B, B’) Different z planes of a germarium show a clone containing an r2a EC, FSCs, and FCs. (C-C’’) Three z planes of a germarium showing a clone with r1 and r2a ECs and FSCs. (D-F) Examples of two clones of different colors in overlapping or adjacent zones, where arrow color indicates the clone color, not the cell type. (D) a GR (GFP plus RFP) clone (yellow arrows) contains r1 ECs and Cap cells (extreme left) and a BR (lacZ plus RFP) clone (magenta arrows) contains r1 and r2a ECs. (E) A BR clone (magenta arrows) contains r1 ECs and a GR clone (yellow arrows) contains a r2a EC and an FSC. (F) A BR clone (magenta arrows) contains ECs and FSCs, and a BG (lacZ GFP) clone (cyan arrows) contains a r2a EC, FSCs and FCs.

**Table 3.** Clone frequencies and cellular yields from different lineage experiments together with deduced single-lineage parameters including the number of precursors of different types.

|  | n | EC |  | EC/FSC |  |  | FSC |  | FC | EC+FC |  | Number of dividing precursors per germarium |  |  |  |  |
| --- | --- | --- | --- | --- | --- | --- | --- | --- | --- | --- | --- | --- | --- | --- | --- | --- |
|  |  | Freq. | Yield | Freq. | Yield |  | Freq. | Yield | Freq. | Freq. | Yield | Total (n) | pEC | pEC/FSC | pFSC | pFC |
|  |  | % | EC | % | EC | FSC | % | FSC | % | % | EC |  |  |  |  |  |
|  |  | 100p |  | 100q |  | x | 100r | y | 100s |  |  | 16/(qx+ry) | pn | qn | rn | sn |
| 0h APFMARCM raw data | 98 | 46.9 | 2.4 | 25 | 4 | 4 | 3 | 4.3 | 9.2 | 16.3 | 2.9 |  |  |  |  |  |
| 0h APF MARCM assuming 70% double clones |  | 63 | 1.8 | 13 | 3.2 | 3.5 | 5 | 4.3 | 19 |  |  | 24 | 15.1 | 3.1 | 1.2 | 4.6 |
| 0h APF Multicolor raw data | 26 | 42.3 | 3 | 31 | 3.5 | 2.6 | 7.7 | 4 | 12 | 7.7 | 2.5 |  |  |  |  |  |
| 0h APF Multicolor assuming 30% double clones |  | 52 | 2.4 | 25 | 2.1 | 2.3 | 9.5 | 3.9 | 14 |  |  | 17.3 | 9 | 4.2 | 1.6 | 2.4 |
| 36h APF MARCM raw data | 121 | 20.7 | 2.5 | 23 | 2.8 | 3.5 | 3.3 | 3.3 | 9.1 | 43.8 | 2.6 |  |  |  |  |  |
| 36h APF MARCM assuming all double clones |  | 51 | 1.8 | 7 | 3 | 4.7 | 4.1 | 3.1 | 38 |  |  | 34.9 | 17.8 | 2.4 | 1.4 | 13 |
| bond-GAL4 raw data | 73 | 38.4 | 5.1 | 48 | 8.3 | 5.4 | 1.4 | 7 | 2.7 | 9.6 | 9 |  |  |  |  |  |
| bond-GAL4 assuming all double clones |  | 61.9 | 3.3 | 24 | 5.6 | 4 | 6.3 | 5.3 | 7.8 |  |  | 15.1 | 9.4 | 3.6 | 1 | 1.2 |

Clones of different colors in the same germarium were generally offset along the AP axis. In some cases, the two differently colored clones likely derive from a single recombination event in a common parent, yielding “twin-spot” daughters (Fig. 3B and Fig. 4D-F). These observations graphically confirm the earlier conclusion that marked lineages spread a limited but significant distance along the AP axis (even for the two daughters of a cell undergoing recombination), frequently spanning boundaries between two different adult cell types.

### EC precursor division declines from the anterior at mid-pupation

The MARCM lineage experiments, where lineages were induced at different times during development, also yield information about when precursors of specific adult cell types stop dividing because only dividing precursors can be labeled by mitotic recombination. For each time of clone induction we summed the total number of marked cells of each type (FSCs, r1 and r2a ECs) over all clones (Table 2). The ratio of marked ECs (r1 or r2a) to FSCs for any one time of clone induction reports the relative number of those cell types produced from dividing precursors. The absolute number of ECs made from dividing precursors was then calculated by using a value of 16 FSCs per germarium and assuming that FSC precursors, like FSCs themselves, do not cease dividing (Table 2). Importantly, this calculation is based on scoring all labeled cells over all clones and is therefore equally valid whether the scored cells in an ovariole derive from a single marked cell or from more than one marked cell.

For −6d and −5d samples, the calculated number of ECs made from dividing precursors was very similar at about 33 (19 r1 and 14 r2a) per germarium (Table 2). This approximates the total number of adult ECs, which has been reported to be about 40 on average [35], and which we counted as 32-51 over 25 germaria from young adults, suggesting that most cells that will become adult ECs are still dividing at the larval/pupal transition. The calculated number of ECs made from dividing precursors dropped slightly in −4d samples (from 33 to 28) and more markedly in −3d (to 15) and −2d samples (to 13) (Table 2). We surmise that precursors of over half of the future adult EC population have ceased dividing by 2-3 days prior to eclosion. Moreover, the average number of dividing precursors for r1 ECs (4.7) was significantly lower than for r2a ECs (8.6) by −2d (72h APF), even though adults contain more r1 than r2a ECs (19 vs 14, from −6d and −5d data). Thus, a reduction in precursor division appears to spread from the anterior towards the posterior as pupal development proceeds.

The inference of few, if any, non-dividing, and therefore potentially fully-differentiated, ECs during the 0-24h APF period is notable because germline cells were found to develop from 1-2 cell units into 16-cell cysts during the same period. These observations suggest that germline progression can be supported by developing EC precursors.

### Some Follicle Cells produced during pupation derive from precursors that also yield FSCs and ECs

In a separate series of lineage experiments, we induced MARCM clones early enough (0h APF and 36h APF) that we were able to examine GFP-labeled cells in newly-eclosed adults (Fig. 3A). This allowed us to score all marked FCs produced from precursors, as well as FSCs and ECs, before the earliest FCs are lost in 2d-old adults. From these data, we can learn about the developmental origin of FCs, which are typically present surrounding the two most mature germarial cysts and four budded egg chambers in a newly-eclosed adult (Fig3A).

We first examined lineages induced 5d before eclosion (at 0h APF). Most ovarioles with marked FSCs also had marked ECs (24/27) (Fig. 5A, D), as observed in MARCM analysis in 2d-old adults and multicolor clone analysis, confirming a common precursor of ECs and FSCs. Almost all clones with marked FSCs also had marked FCs (25/27) (Fig. 5A), indicating a common precursor for adult FSCs and FCs. A significant proportion of all marked ovarioles (23/98=23%) had marked ECs, FSCs and FCs, indicating that some individual pupal precursors give rise to all three cell types in newly-eclosed adults. Many ovarioles (“FC-only”) contained marked FCs but no marked FSCs (Fig. 5C, D). Similar results were observed for multicolor lineage tracing examined in newly-eclosed adults (Fig. 4, Table 3). These results suggest that at the start of pupation there is a group of anterior precursors that produces ECs, a group of posterior precursors that produces only FCs and a group of precursors in between that produce a combination of two or three of these cell types (Fig. 6).

**Figure 5.**
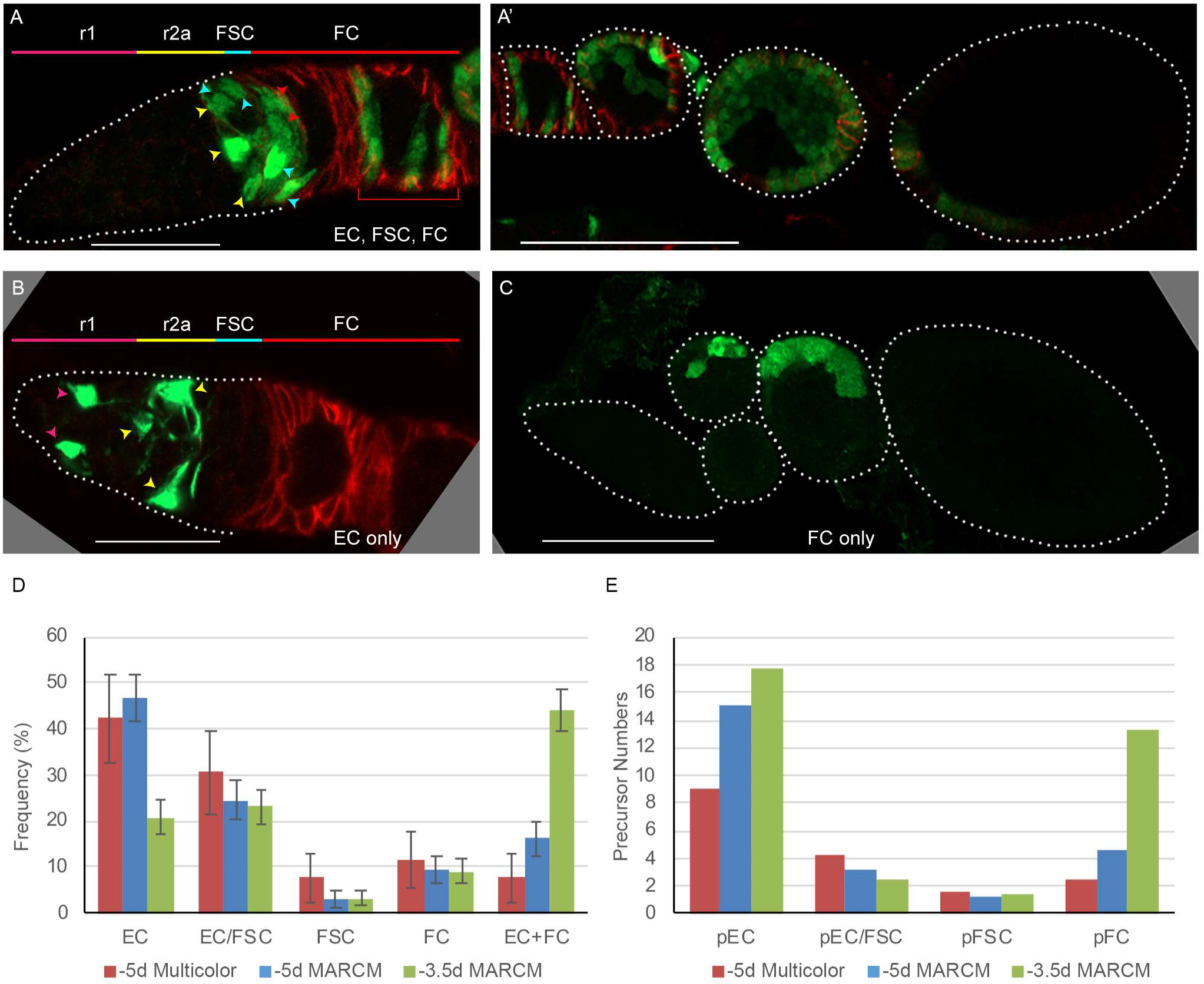
Follicle Cell origins and deduced precursor numbers for ECs, FSCs and FCs. (A-C) GFP-labeled MARCM clones (green) were induced 5d before eclosion and ovaries were dissected from newly-eclosed adults in order to detect all FCs produced during pupation. Fas3 is stained in red and cell types are indicated with color-coded arrows, matching the labeled AP domains. Scale bar in (A, B), 20 μm; (A’, C), 50 μm. (A) Germarium and (A’) the rest of the ovariole of the same sample (the first egg chamber is in both images), showing a clone with marked ECs, FSCs and FCs (marked FCs are in the germarium and egg chambers 1-4). (B) A clone containing only ECs (no cells were marked in the rest of the ovariole). (C) A clone containing only FCs in egg chambers 2 and 3. (D) Distribution of clone types in GFP-only clones of multicolor flies and MARCM clones initiated 5d or 3.5d before eclosion, all examined in newly-eclosed adults. Clone types are: EC (EC-only), EC/FSC (ECs and FSCs, with or without FCs), FSC (with or without FCs), FC (FC-only), and EC + FC. Ovarioles with only marked ECs and FCs likely harbor two independent lineages. (E) Estimates of numbers of dividing precursor types at the time of clone induction for the same three lineage experiments as in (D), after accounting for double-clones (see Methods and Table 3). Precursor types are defined by their adult products: pEC (only ECs), pEC/FSC (ECs and FSCs, with or without FCs), pFSC (FSCs with or without FCs), and pFC (only FCs).

**Figure 6.**
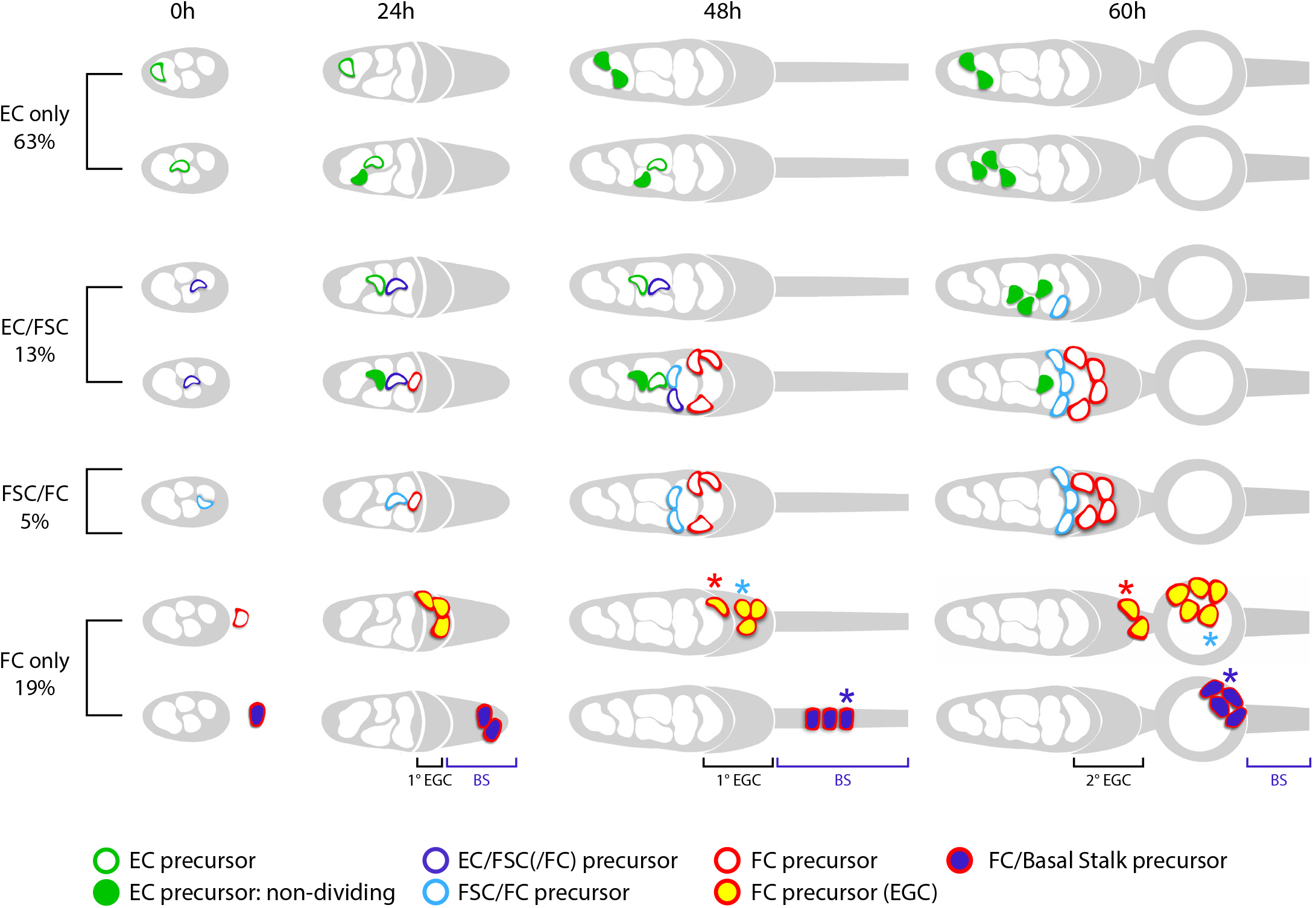
Summary of precursor lineage development. Cartoon of the fate of single precursor types from 0h APF until 60h APF deduced from lineage studies and pupal ovary imaging. The indicated frequency of clone types is drawn from the inference of single-cell lineages for MARCM clones induced at 0h APF (Table 3). Most 0h APF precursors (green) generate only adult ECs, with more anterior (left) precursors producing only r1 ECs (top row) and more posterior precursors also producing r2a ECs and more cells in total (second row). Filled green cells have ceased division. Most 0h APF FSC-producing precursors also produce ECs. Some of the more anterior precursors (third row) do not produce FCs but most (fourth row) produce cells that will become FCs (but not in the first-formed egg chamber). Precursors that produce adult FSCs but not ECs also produce FCs (fifth row). The most posterior precursors produce only FCs. Examples of origination from an EGC (row 6) and a basal stalk cells (row 7) are shown, but the most posterior ICs can also yield lineages containing only FCs. A subset of progeny accumulate posterior to the germarium from 24h APF onward in the EGC (depicted as the posterior gray crown). Most cells in the EGC will become FCs of the first budded egg chamber (sixth row). The cell on the posterior cyst at 48h APF, denoted by the red asterisk, moved into the secondary EGC at 60h and is modeled on the top red cell in Fig. 10. The cell denoted by the blue asterisk that is in the EGC at 48h APF and moved onto the first budded cyst at 60h is modeled after the cyan cell in Fig. 10. Most precursors present in the basal stalk at 48h APF (row 7) move onto the first budded cyst (the cell denoted by the blue asterisk is modeled after the white cell in Fig. 10). More posterior precursors are shown producing larger numbers of progeny because anterior cells divide more slowly and arrest division prior to eclosion.

The locations of marked FCs were scored as “immediate” (region 2b), “germarial” (region 3) or in egg chambers (“ECh”) 1-4 (Fig. 3A and Fig. 7A). The percentage of clones with marked immediate FCs was higher for ovarioles that included marked FSCs (56%) than for those without marked FSCs (16%). Conversely, the percentage of ovarioles with marked FCs in at least one of the two terminal (most posterior) egg chambers was higher when marked FSCs were absent (88%) than when they were present (67%). Indeed, occupancy by marked FCs decreased gradually from anterior to posterior for FSC/FC clones and with the opposite polarity for FC-only clones (Fig. 7A). Thus, derivatives of a given precursor were clustered along the AP axis in FSC/FC territory, as previously noted for EC/FSC territory. The inference that the fate of precursor derivatives is set largely by the AP position of precursors, with limited AP dispersal of derivatives, appears to be the governing principle all the way from the most anterior r1 EC derivatives to the most posterior FCs in newly-eclosed adults. Notably, while some clones contained only ECs or only FCs, the AP spread of derivatives generally exceeded the width of the FSC domain, so that ovarioles with marked FSCs always contained ECs or FCs (or both). In other words, there were no FSC-specific precursors at the larval/pupal transition and individual lineages regularly spanned the borders of adult cell identities (Fig. 6).

**Figure 7.**
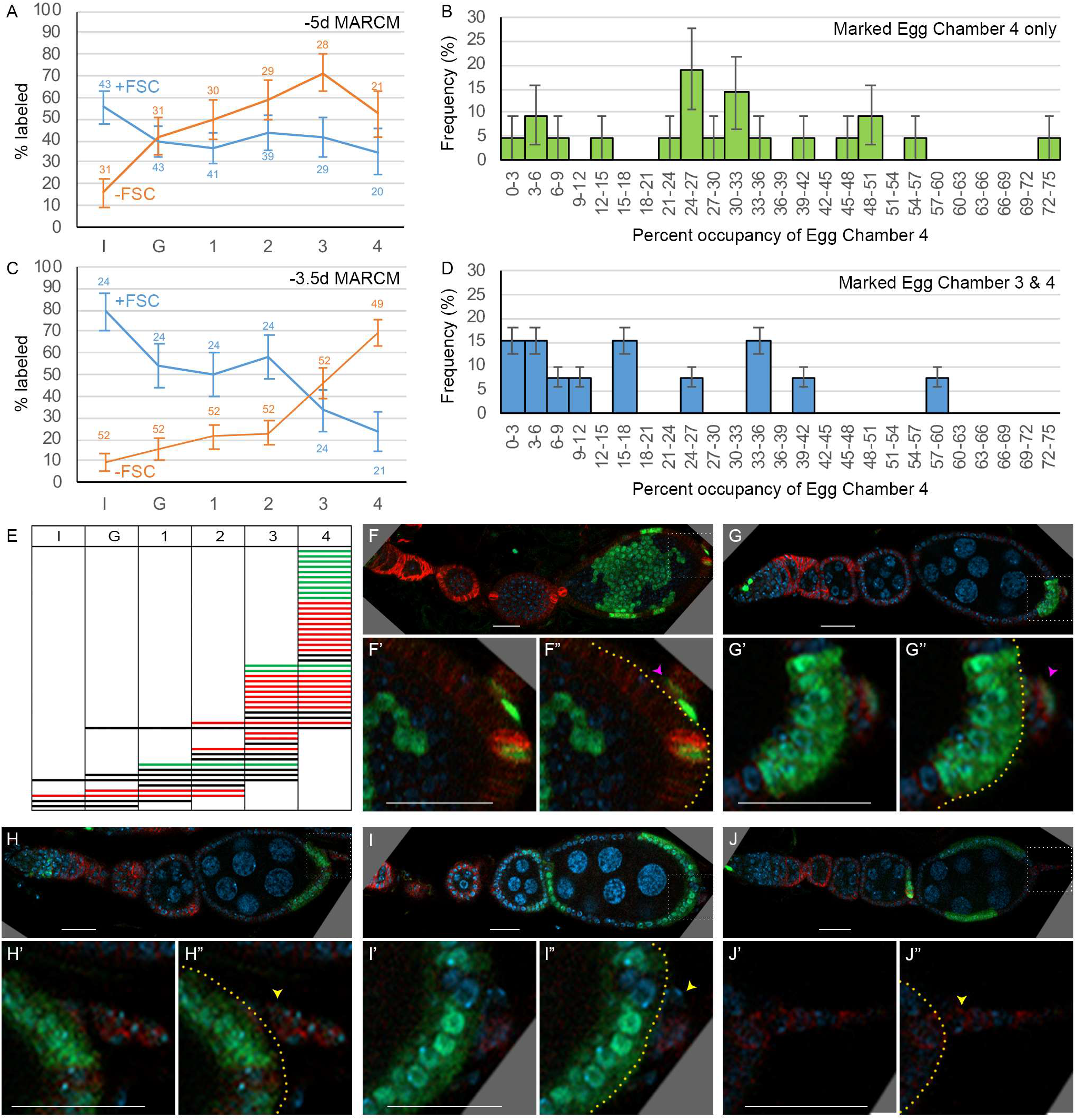
Patterns of FC contributions indicate distinct pools of precursors along the AP axis. (A, C) Percentage of ovarioles containing labeled FCs that have marked FCs in germarial region 2b (I), region 3 (G), or egg chambers 1, 2, 3 or 4 (see Fig. 3A) in newly-eclosed adults for (A) −5d MARCM and (C) −3.5d MARCM clones with FSCs (blue line) or with no FSCs (orange line). The total number of ovarioles scored is indicated either side of the error bars (SEM); some ovarioles had only 3 egg chambers and some were not fully imaged. (B, D) MARCM clones induced at −3.5d and analyzed in newly-eclosed adults. Marked occupancy of egg chambers was scored as a percentage of all FCs in egg chamber 4 that were marked when there were marked FCs (B) only in Egg Chamber 4 or (D) in egg chambers 3 and 4. The frequency of different occupancy ranges is displayed in bins of 3%. (E) Distribution amongst germarial region 2b (I), region 3 (G) and egg chambers 1-4 of FC-only clones in newly-eclosed flies after marking at −3.5d. Each row indicates a single ovariole. The colors indicate if a marked cell is present in the basal stalk (green), absent from the basal stalk (red) or if the basal stalk was not clearly imaged (black). (F-J) Ovarioles of newly-eclosed flies with MARCM recombination induced at −3.5d containing FC-only clones. (F, G) Marked FCs can be seen in the basal stalk and egg chamber 4, (H) egg chamber 4 only, or (I, J) in egg chambers 4 and 3 only. Insets show enlargements of boxed regions; yellow borders outline the posterior edge of Egg Chamber 4, with cells further posterior located in the basal stalk. Yellow arrowheads (no marked basal stalk cell); Pink arrowheads (marked basal stalk cell). GFP: green; Fas3: red; DAPI: blue. Scale bars, 20 µm.

### Estimates of numbers of different precursors just prior to pupation

The number of precursors at 0h APF that give rise to different adult cell types can be deduced from the observed proportion of lineage categories (EC, EC/FSC+/-FC, FSC+/-FC, FC) together with the average number of marked FSCs and ECs in each lineage category. However, both measured parameters will be inaccurate if some ovarioles include two or more marked lineages. Analysis of lineages in 0d adults allowed us to estimate how often this occurred.

In this MARCM experiment, 67% of 98 ovarioles contained no marked cells, suggesting that 80% of marked ovarioles have lineages derived from a single cell if all recombination events are independent (see Methods). However, 16% of marked ovarioles contained marked FCs and marked ECs but no marked FSCs, strongly suggesting that these discontinuous patterns represent two lineages and hence, that the total proportion of ovarioles with more than one marked lineage is considerably higher than 20%. Thus, there appears to be some clustering of recombination events rather than an even distribution among all ovarioles, perhaps because temperature changes are not uniform during the mild heat-shock conditions used to induce recombination events at a very low frequency. The observed proportions of ovarioles with marked ECs and FCs roughly fit a simplified model where 30% of ovarioles with any marked cells derive from a single marked cell and 70% derive from exactly two marked cells (see Methods). We used that model to calculate the component single cell lineage frequencies and component cell numbers per single-cell lineage that would give rise to the scored raw data (see Methods). The deduced average number of dividing precursors at pupariation was about 24 in total, with 15.1 precursors giving rise only to ECs, 3.1 precursors producing ECs and FSCs, 1.2 precursors producing FSCs but no ECs, and 4.6 precursors producing only FCs on average (Fig. 5E; Table 3).

Multicolor GFP-only clones were used previously to provide the most definitive evidence that an individual precursor cell can produce both adult ECs and FSCs. The frequency of ovarioles with marked ECs and FCs but no FSCs was only 8%, consistent with a high proportion of single cell lineages, which we estimated at 70% (see Methods). From the inferred proportions of single lineage types and their cellular contents (see Methods), we deduced that a single precursor cell gives rise to both ECs and FSCs in 25% of all marked lineages and that the frequency of such precursors greatly exceeds precursors that produce FSCs but no ECs (10%). The deduced proportions of clone types and the number of precursors of each type (9.0 EC, 4.2 EC/FSC, 1.6 FSC, 2.4 FC and 17 total precursors) are reasonably similar to those calculated above from the similarly timed MARCM experiment (Fig. 5E; Table 3). Differences may result from variations in the exact developmental stage of animals of different genotypes at the time of clone induction.

We also performed a MARCM analysis of clones induced 3.5d prior to eclosion (36h APF) and examined in 0d adults to learn more about the timing of FC specification (see below). In this experiment, the proportion of ovarioles that were labeled (94%) and the frequency of EC/FC clones was particularly high, probably because there are many more dividing precursors than at 0h APF, so the deduction of single cell lineage content is likely less reliable. Nevertheless, we estimated (see Methods) that the proportions of single-cell lineage types were EC-only (51%), EC/FSC(+/-FC) (7%), FSC(+/-FC) (4%), FC-only (38%) and that the total number of dividing precursors was about 35 (Fig. 5E; Table 3). The most striking difference between lineages induced at 36h versus 0h APF is a large increase in lineages producing only FCs. Based on earlier analysis of MARCM lineages induced at −4d and −3d and examined at +2d, there should also be between 5 and 18 non-dividing EC precursors at −3.5d (Table 2). Thus, we estimate that there are 13.3 FC-only precursors within a total precursor population of about 40-53 (35 dividing plus 5-18 non-dividing cells) at 36h APF (Table 3).

### Timing of FC recruitment to first egg chamber

To determine when FCs are first allocated to the most mature germline cyst, we looked at the distribution of marked cells in FC-only clones. The majority of newly-eclosed adults have four egg chambers, and we excluded all exceptions from the analysis described below. For FC-only clones induced at 36h APF, more than half (34 out of 49) included labeled FCs in the fourth (most mature) egg chamber. Two-thirds of those (21 out of 34) had no other labeled FCs, showing that many precursors are restricted by 36h APF to contribute only to the first-formed egg chamber (Fig. 7E, F). The 13 exceptions all had marked FCs in egg chamber 3 (Fig. 7E, G). This frequency is too high for all to be explained by the presence of two lineages, and a single lineage might plausibly contribute to two adjacent egg chambers. We also estimated the proportion of the egg chamber 4 epithelium occupied by marked FCs. The proportion was significantly lower when marked FCs were also in egg chamber 3 (19.6%) than when confined to only the terminal egg chamber (29.7%) (Fig. 7B, D). These results suggest that, in most cases, the initially labeled precursor at 36h APF divided before all egg chamber 4 FCs were allocated. In some cases, one daughter became an FC founder for egg chamber 4 and the other daughter later contributed to egg chamber 3. In other cases, both daughters contributed to egg chamber 4, accounting for a higher average FC contribution for that labeling pattern. We do not know the cell cycle time for these FC-only precursors subsequent to 36h APF, but we expect (from amplification data for EC/FSC lineages in Table 1) that it is shorter than 24h and may be as short as 11h, the cycling time for early FCs in adult ovarioles [68], so we estimate that FCs are allocated to egg chamber 4 at about 47-60h APF. This estimate fits with later, direct morphological observations.

By 48h APF, all germaria have a single cyst in the most posterior region (Fig. 2) but neighboring somatic cells (ICs) only express very low levels of Fas3 (Fig. 1G). In adult germaria, even the most recently recruited FCs surrounding a stage 2b cyst express high levels of Fas3; a subset of those cells must then become specialized polar and stalk cells before the egg chamber can bud a further 24h later (Fig. 8J). The most posterior ICs that do not express Fas3 strongly before 48h APF therefore appear to be far behind schedule, given that the first egg chamber will bud at about 56h APF on average (Fig. 1). These observations raise the question of whether these ICs are indeed the cells that will form the FCs of the first budded egg chamber, as might have been assumed by analogy to budding of adult egg chambers.

**Figure 8.**
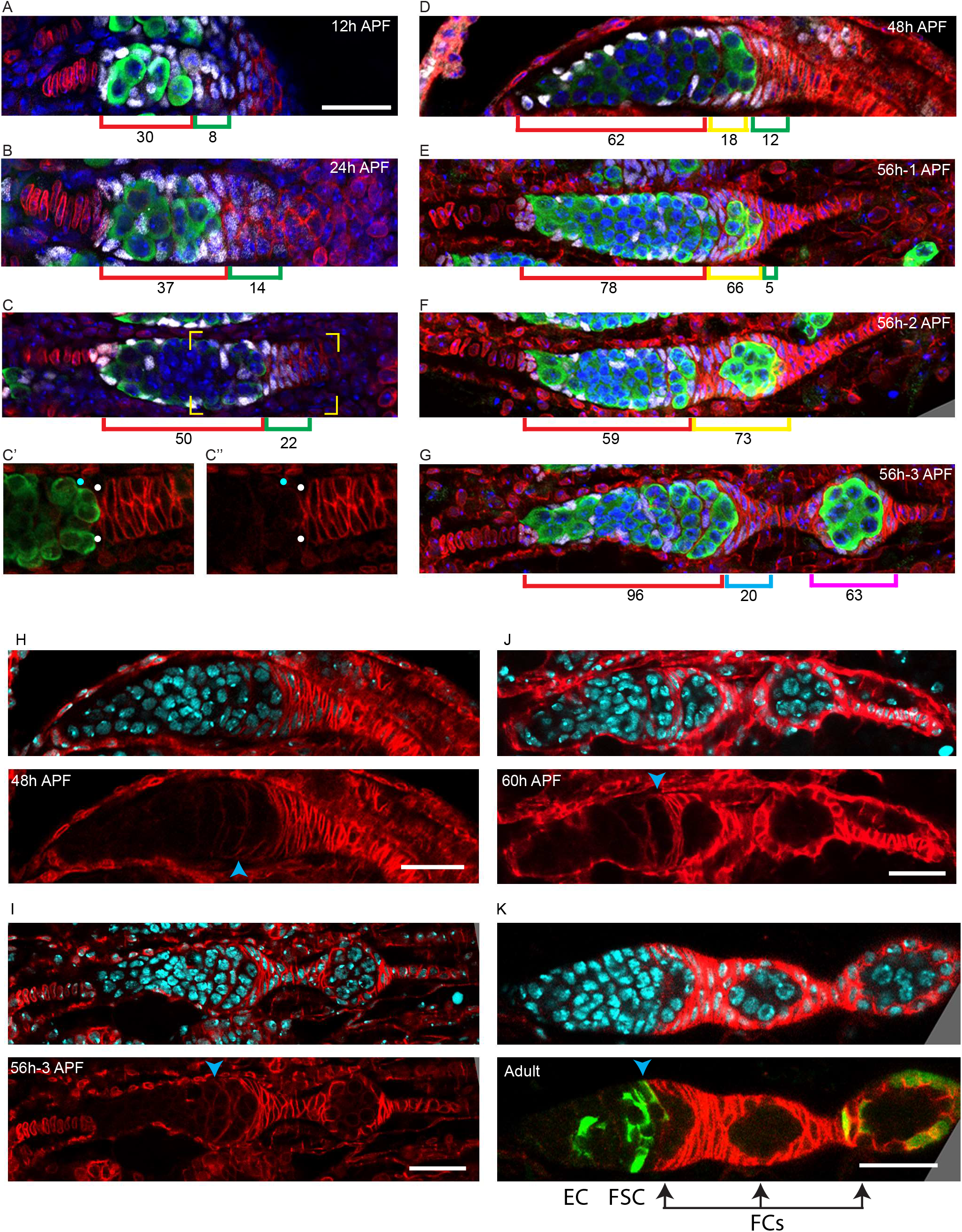
Extra-Germarial Crown Cells (EGCs); a potential source of early FCs. (A-G) Traffic Jam (TJ) positive cells (white) were counted in ovaries stained also for Vasa (green), LaminC and Fas3 (both in red), and DAPI (blue). Bar, 20 μm applies to all images. (A-C) Up to 36h APF, prior to budding of the first egg chamber, ICs (red brackets), defined as intermingled with germline cells (green) in the developing germarium, expressed TJ (white) but little or no detectable Fas3 (red). Cells posterior to the developing germaria expressed Fas3 and were previously all termed basal cells and expected to intercalate to form the basal stalk. However, a subset of Fas3-positive cells immediately posterior to the germarium also expressed TJ. Those cells formed an Extra-Germarial Crown (EGC) of increasing cell numbers, indicated by green brackets. (C’, C”) At 36h APF some somatic cells contacting the posterior of the most advanced cyst expressed Fas3 strongly (white dots) and some did not (blue dots). Fas3-positive cells were counted as EGC cells and Fas3-negative cells as ICs. (D) At 48h APF the most posterior germline cyst is about to leave the germarium in most samples. Here, the cyst (green) appears to have just started to leave, invading Fas3-positive cells in EGC territory. The Fas3-positive cells surrounding the cyst are indicated by a yellow bracket, while Fas3-positive and TJ-positive cells posterior to the cyst are indicated by green brackets. Average IC (red brackets) and EGC (green and yellow brackets) cell numbers are written underneath the brackets and were derived from counting (A) 5, (B) 5, (C) 10 and (D) 5 samples. (E-G) Three developing ovarioles from the same 56h APF ovary, representing progressively later stages of development from top to bottom, show the most posterior cyst leaving the germarium and surrounded by Fas3-positive, TJ-positive cells. (E, F) Fas3-positive, TJ-positive cells posterior to the germline (green bracket), and around the most posterior cyst (yellow bracket) were counted for these single samples and then (G) counted as on the budded cyst (magenta bracket) or anterior to the budded cyst and forming a secondary EGC posterior to the germarium (blue bracket). Those two populations are separated by stalk cells that do not stain for TJ. (H-K) Fas3 (red) is shown together with DAPI-stained nuclei (blue; top) or alone (bottom) in pupal ovaries at (H) 48h APF, (I) 56h APF (same samples as in (D) and (G)), (J) 60h APF and (K) adults. The anterior border of strong Fas3 expression (indicated by a blue arrowhead) was (C’, C”) posterior to all germline cysts at 36h APF, aligns (H) with the anterior of the departing cyst at 48h APF, (J, K) with the anterior surface of the most posterior germarial cyst just after one egg chamber has budded, and (J) by 60h APF and (K) in adults, the anterior edge of strong Fas3 expression is along the posterior edge of the penultimate cyst in the germarium. (K) Cells in a MARCM lineage (green) derived from an adult FSC (bottom panel) include an EC, FSCs and FCs (outlined by strong Fas3) in the budded egg chamber. Arrows indicate the normal progression every 12h of a recently produced strongly Fas3-positive FC associated with the penultimate germarial (stage 2b) cyst (first arrow) to an FC on a stage 3 germline cyst at the posterior of the germarium (second arrow) and then to an FC on a budded egg chamber (third arrow); 24h in total.

### Lineage evidence for separate precursors of FCs in the first-formed egg chamber

Marked FCs occupied only the two terminal egg chambers (one or both) in 35 out of 49 ovarioles with FC-only clones induced at 36h APF (Fig. 7E). Marked FCs occupied only territory anterior to the two terminal egg chambers in five ovarioles, while the remaining nine ovarioles included labeled FCs in terminal egg chambers and more anterior locations. It was estimated that most ovarioles in this experiment have two marked lineages and that 38% of all single-cell lineages contain only FCs, so around 11.5 of the 49 FC-only ovarioles (49 x (0.38)^2^/[(2 x 0.38 x 0.62) + (0.38)^2^]) might be expected to include two FC-only lineages. Importantly, there were only two ovarioles with labeled FCs in egg chamber 4 and also in locations anterior to egg chamber 3 (Fig. 6E) and these may plausibly contain two lineages, consistent with a conclusion that, at 36h APF, precursors to FCs of the first-formed egg chamber contribute only to egg chambers 3 and 4 (seen in 32/34 cases).

It is possible that the seven ovarioles with labeled FCs in egg chamber 3 (but not 4) and in more anterior locations all harbor two lineages, but that seems unlikely since all seven have marked FCs in egg chamber 2 (suggesting a common origin) and there were only 23 ovarioles with marked FCs in egg chamber 3 altogether. Thus, by 36h APF the FC-only precursors appear to be segregated into two populations. The majority contribute FCs to the first-formed egg chamber and sometimes also to egg chamber 3, while the remainder contribute FCs to more anterior egg chambers, sometimes including egg chamber 3. The AP location of different FC-only precursors at 36h APF presumably accounts for these differences.

Ovarioles that included marked FSCs only rarely had marked FCs in the terminal egg chamber (Fig. 6C). The five exceptions (out of 21) may derive from ovarioles that harbor an FC-only lineage (53% frequency overall) in addition to an FSC-containing lineage (26% overall). Thus, at 36h APF it appears that around two-thirds of FC-only precursors will contribute to the first-formed egg chamber, with some of these also contributing to egg chamber 3, while more anterior egg chambers are later populated by derivatives of a combination of FC-only, FSC/FC and EC/FSC/FC precursors.

### An Extra-Germarial Crown (EGC) forms prior to egg chamber budding

Our cell lineage studies, in which any dividing cell can be labeled, provided information about gradual restriction of precursor fates during pupation, according to cell locations along the AP axis. To define the location of the precursors of FCs that coat the first budded egg chamber, we first examined the changing morphology and gene expression patterns of developing ovaries. We already described the onset of weak Fas3 expression in ICs as surprisingly close to the time of egg chamber budding (Fig. 1), suggesting that ICs may not be the source of the first FCs because the first FCs most likely already express Fas3 strongly.

Posterior to the interspersed germline cells and somatic ICs of the pupal germarium are somatic cells with strong Fas3 staining from 24h APF onward (Fig. 1 and Fig. 8). We noticed that the most anterior of these Fas3-positive cells express TJ, which is also expressed in ICs and required for their interspersion with germline cells [69]. The TJ-positive cells form a small multi-layered cone or crown around the posterior of the developing germarium that increases in size and cellular content from 12-36h APF (Fig. 8A-C). We refer to those TJ/Fas3-positive cells beyond the most posterior germline cyst as the Extra-Germarial Crown (EGC). A population of cells that expresses both TJ and Fas3 strongly is notable because there is evidence from manipulation of TJ activity in adult ovaries that TJ generally represses Fas3 expression [69], while a principal characteristic that distinguishes adult FCs from other TJ-expressing cells (FSCs and ECs) is strong Fas3 expression (Fig. 8K) [23]. What becomes of the newly-identified group of EGC cells, which express Fas3 strongly in addition to TJ?

In adults, a stage 3 germline cyst is separated from the most recently budded egg chamber by only a small number of somatic cells (Fig. 1J and 8L). The stage 3 cyst then buds from the germarium together with adjacent somatic FCs, which emerge as a monolayer epithelium connected to the germarium by a thin, short stalk of specialized follicle cells. By contrast, prior to budding of the first pupal egg chamber there is a large accumulation of somatic EGC cells posterior to the germarium (Fig. 8A-D) and after budding, the first egg chamber is often found much further from the germarium than in adults (Fig. 1H). The intervening space is occupied by a partially intercalated stalk of TJ-negative cells and by a small “secondary” EGC of Fas3- and TJ-positive cells, crowning the germarium (Fig. 8G). After one cyst has budded and a second cyst has rounded up in preparation for budding (around 60h APF) there is strong Fas3 expression in germarial somatic cells (Fig. 8J), resembling the pattern seen in adults (Fig. 8K). In samples with two budded egg chambers (72-84h APF) or more, the newly budded egg chamber is close to the germarium (Fig. 1I). Thus, formation of the first egg chamber appears to involve a morphological process quite different from adult egg chamber budding; this is partly recapitulated by the second egg chamber before adopting a typical adult morphology.

### Live imaging reveals EGC and basal stalk cells as the source of FCs for the first egg chamber

We used a live imaging protocol that we had previously developed for adult ovarioles [70] in order to view pupal ovaries and track individual somatic cells. We were able to follow isolated live ovaries for up to 15h, during which time many cell movements and cell divisions were observed, suggestive of continued normal behavior. We first used samples where all cells were labeled by expression of a *ubi-GFP* transgene. Later, we used animals with multicolor clones induced 2-3d earlier, so that some cells had lost GFP or RFP expression or both (Fig 3B). Variability in the levels of GFP and RFP simplified the task of following individual cells over time. We were able to track the movements of multiple cells in a series of videos initiated at a variety of times after pupariation.

From about 30h APF onward, tracked ICs moved short distances relative to other cells in a variety of directions. Most ICs maintained their relative AP locations within the developing germarium as cysts moved past them. For example, one of two initially parallel germline cysts adopted the most posterior position in the germarium while adjacent somatic ICs remained in position (Fig. 9A, B). Even cells posterior to the most posterior cyst at 40h (red arrow, Fig. 9C), which are likely EGC cells (Fig. 8C), moved relative to the cyst to end up at the anterior of the cyst several hours later. At the same time, an EGC cell (magenta arrow, Fig. 9C), initially distant from the cyst was adjacent to the cyst 9h later. Another 40h APF movie similarly showed the cyst moving away from ICs (yellow, cyan, white, magenta fill, in Fig. 9B) and making contact with EGC cells (outlined red cells, Fig. 9B). These observations support the deductions from lineage analysis that precursors undergo limited, stochastic movements along the AP axis and suggest that 16-cell germline cysts do not have a stable coating of somatic cells from about 30-50h APF. Instead, by 48h APF the most posterior germline cyst is starting to move into EGC territory. These observations are consistent with fixed images showing Fas3-positive cells only posterior to the cyst at 36h APF (Fig. 8C-C”) but beginning to surround the cyst by 48h APF (Fig. 8D) because the cyst moves into the Fas3-positive EGC domain. During the 30-50h APF period, there were cell divisions in the EGC and stalk (Movie S1) but we saw no evidence of cells moving from the IC region into the EGC or in the reverse direction (see outlined red EGC cells in Fig. 9B), suggesting that the EGC population grows primarily as a result of EGC cell division.

**Figure 9.**
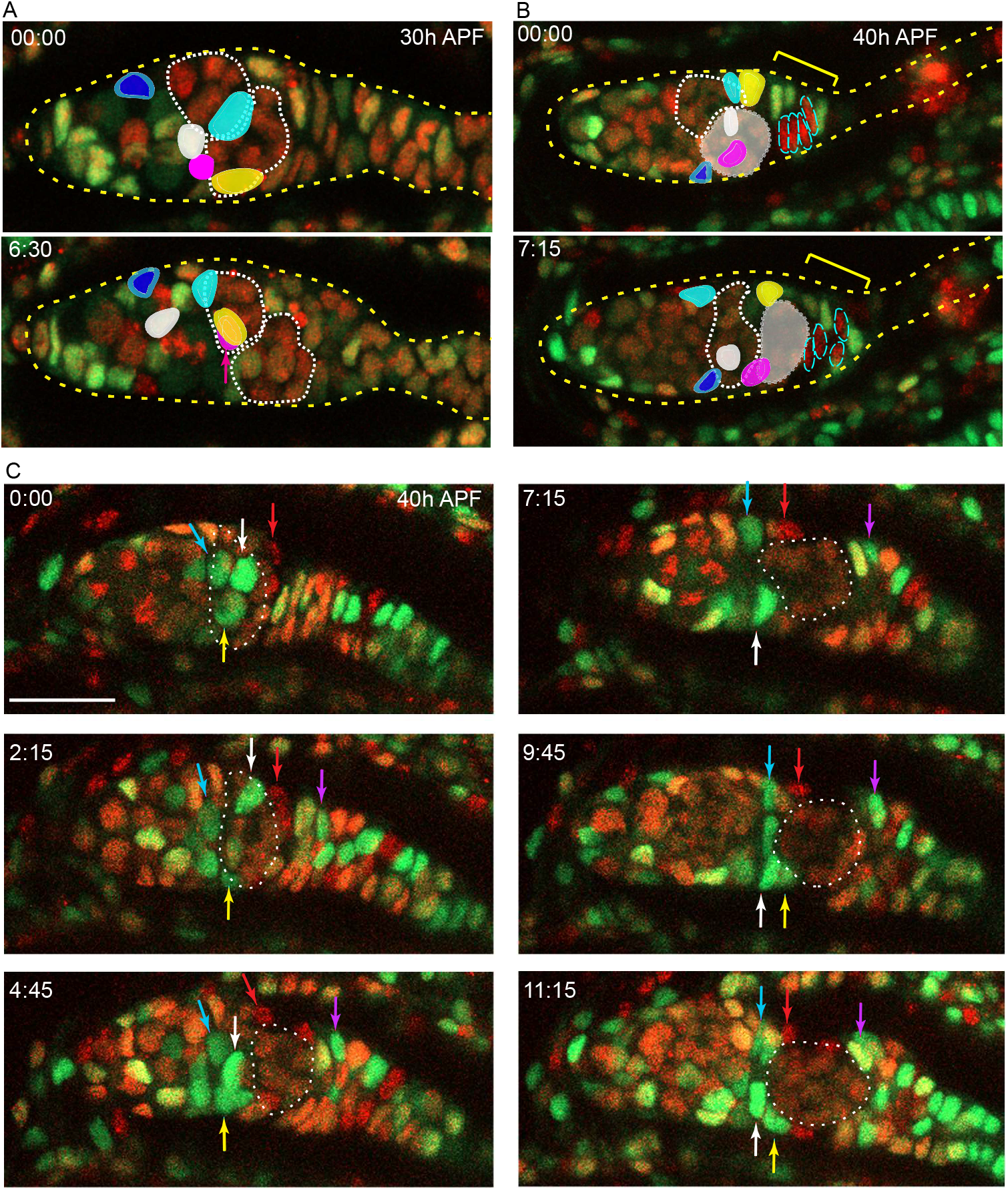
ICs move independently of one another and do not mix with EGCs. Live imaging was performed on multicolor pupal ovaries with clones induced by heat shock such that cells could lose RFP, GFP, or both. (A) Z-projections of 0h and 6h30min timepoints of a 30h-APF germarium showing five tracked ICs (colored). ICs were located at the outer surface of the germarium and moved independently of one another in posterior, anterior, and lateral directions (around the circumference). The most posterior germline cyst (outlined by a white dotted line) moved posteriorly past the cyan and yellow IC cells. The relative disposition of ICs and cysts changed considerably, suggesting they are not strongly associated. Only z planes containing tracked cells were included in the projection (z4, 5, 6, 7, and 8 at 00:00 and z5, 6, 9, and 10 at 6:30) (B) Z-projections of 0h and 7h15min timepoints of a 40h-APF germarium showing five tracked ICs. ICs moved independently of one another and remained in their domain as the most posterior germline cyst (highlighted by shading) moved past. Four cells in the EGC (outlined red cells) remained in the EGC (indicated by the yellow bracket). Projected sections are z5-9 at 00:00 and z4, 7, 8 and 9 at 7:15. Individual z sections of the germaria in A and B are shown in Fig. S2. (C) Z-projections of selected timepoints of a 40h-APF germarium imaged for 11h15min, showing three tracked green ICs (white, yellow and blue arrows); ICs remained in their domain as the posterior germline cyst (outlined by a white dotted line) moved past them. Four red cells (one is shown by the red arrow) started on the posterior of the posterior cyst and moved anterior along the cyst; cells in a neighboring no-color clone moved similarly. A green cell that began in the EGC (magenta arrow) moved onto the posterior edge of the cyst. The accompanying Movie S1 of this germarium shows mitotic cells in the EGC and the stalk. Bar, 20 µm.

Imaging of an ovary from about 48h APF onwards captured the key process of egg chamber budding for the first time (Fig. 10; Fig. S3-S5; Movie S2-S4). At 48h APF, fixed images showed that cells surrounding the posterior half of the germline cyst express Fas3 strongly (Fig. 8D, yellow bracket), consistent with evidence from live imaging that the cyst is already moving into EGC territory (Fig. 9B, C), as described above. The location of Fas3-positive TJ-positive EGC cells is inferred (yellow and green brackets, Fig. 10) by comparison to similarly-staged fixed images (Fig. 8), with Fas3-positive TJ-negative cells posterior to the EGC considered basal stalk cells. The most posterior germline cyst (highlighted by green nuclei in the lower germarium of Fig. 10) moved out of the germarium into EGC territory, as EGC cells enveloped the cyst. For example, the cyan-outlined cell started within the EGC, contacted the cyst prior to 8.3h and remained associated for the next 6.5h (Fig. 10; Fig. S4). Similarly, the pink cell, initially just posterior to the cyst, moved to the midpoint of the budded egg chamber (Fig. 10; Fig. S3C; Movie S2). The red-outlined cell (red asterisk), which started on the posterior face of the cyst, like the pink cell, initially maintained contact with the posterior-moving cyst. It was last seen at, or just beyond, the anterior aspect of that cyst, suggesting it is moving past the cyst to contribute to the secondary EGC, awaiting the second cyst to bud (Fig. 10; Movie S3).

**Figure 10.**
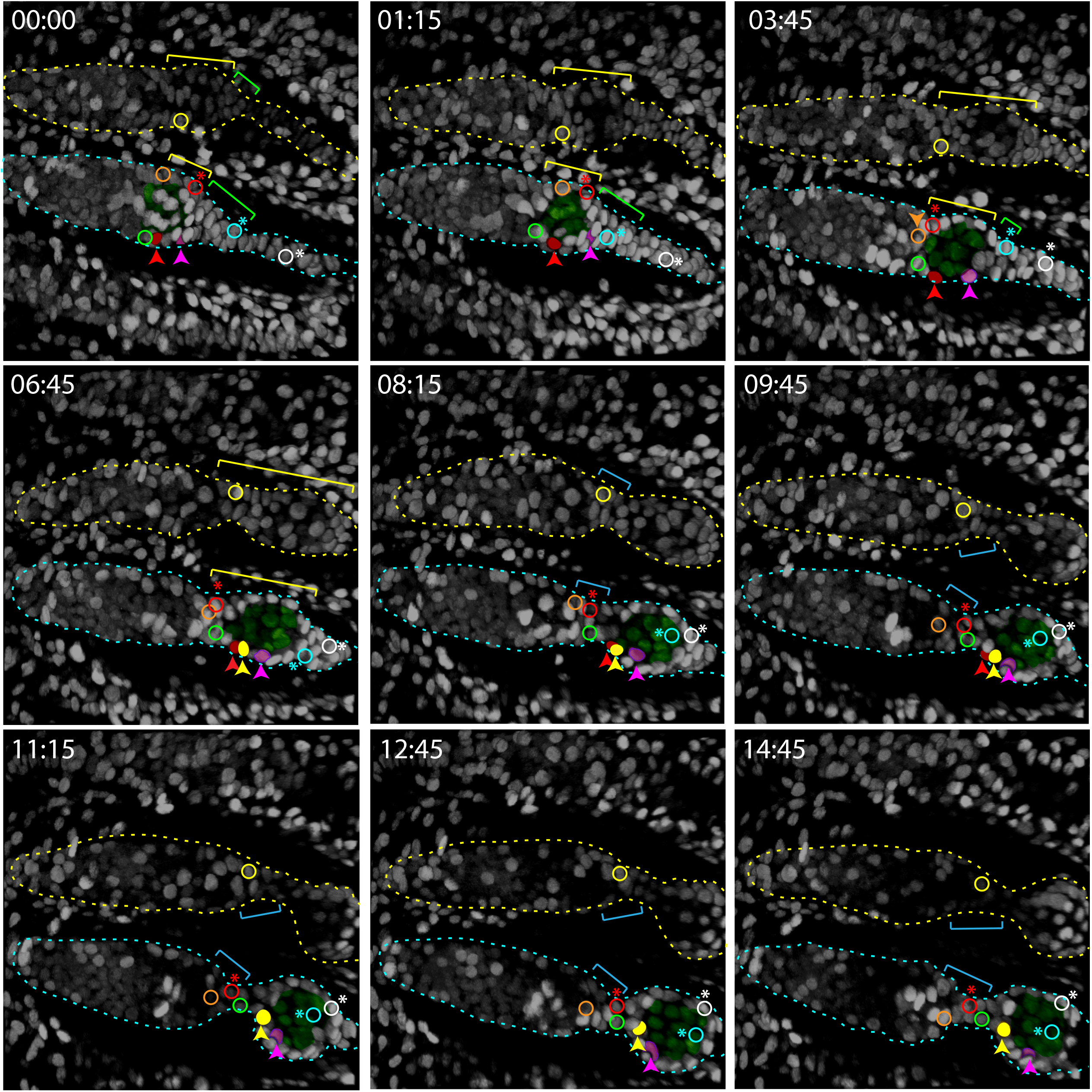
Budding of the first egg chamber: EGC cells become FCs. Time-stamped frames from a 3D reconstruction of a movie of 14h 45 min starting at 48h APF and showing budding of an egg chamber. All cells are labeled white from a ubi-GFP transgene. Three cells that were visible near the surface of a 3D reconstruction are colored in solid red, pink, and yellow in the lower germarium, together with five other cells (open circles) that were in interior z slices. All marked cells were tracked using Zeiss Zen software for 60 time points over 14h 45 min, except for the red cell which could not be reliably tracked after 10h and the yellow cell in the lower germarium, which was not tracked before 5h 45min. The posterior germline cyst in the lower germarium is highlighted in green. Cyst cells were distinguishable from somatic cells because the nuclei were larger and paler green. The yellow brackets indicate Fas3+ TJ+ EGC cells, inferred from comparison to fixed images at the same stages. The cyst slid past adjacent ICs, which were later either at the posterior edge of the germarium (orange-outlined cell and yellow-outlined cell in the upper germarium) or between the germarium and the budded cyst (green-outlined cell) in the nascent secondary EGC (magenta brackets), as seen more clearly in Fig. S2A, B (yellow-outlined cell) and Fig. S4A, B (orange and green-outlined cells). The cell marked with the open red circle (asterisk; modeled in Fig. 6) is inferred from comparison to fixed 48h APF images to be Fas3-positive and to be one of the most anterior EGC cells when the cyst first started to move out of the germarium (just prior to 48h APF, see Fig. 9). This cell began on the posterior cyst and ended in the secondary EGC (Movie S3). The cyan cell (asterisk; modeled in Fig. 6) initially towards the posterior of the primary EGC (indicated by yellow brackets) adopted progressively more anterior positions on the germline cyst after the cyst moved into EGC territory. The white cell (asterisk; modeled in Fig. 6) was originally in a location posterior to the EGC (it is expected, from comparison to fixed images, not to express TJ) but then moved into EGC territory by 3h 45min and ended up close to the cyan cell on the budded germline cyst (Movie S4). The locations of the cyan and white cells covering the posterior of the cyst can be seen clearly in Fig. S4. Individual z-sections showing the locations of the red, pink and yellow cells in the lower germarium at the beginning, middle, and end of their imaging times are shown in Fig.S3C and Movie S2. Time is in hours:minutes.

IC cells were largely left behind in the germarium as the germline cyst moved into EGC territory. The cells outlined in orange and green, like the yellow-outlined cell in the germarium above, started close to the anterior face of the terminal cyst and lost contact with the cyst before 7h. At the end of the imaging period, the cells outlined in orange and yellow were adjacent to the second cyst, inside the germarium (Fig. 10; Fig. S3B and S5B). The green-outlined cell at 14:45h is in a location likely to become the secondary EGC. The secondary EGC may therefore result from cells that were originally ICs (green-outlined cell) and in the EGC (red asterisk).

Surprisingly, we observed that even distant basal stalk cells migrated onto the budding germline cyst. The white-outlined cell, which initially lay posterior to the region with TJ expression (from comparison to fixed images), entered EGC territory within 4h and became associated with the germline cyst by 12h (Fig. 10; Fig. S4 and Movie S4). Thus, over the nearly 15h of live imaging the newly-budded cyst acquired a set of closely-associated cells of epithelial appearance that derived from the EGC and basal stalk cells, while tracked ICs at 48h APF were anterior to the budded cyst 15h later.

Live imaging of a different ovariole from 55h APF onward showed continued posterior movement of the budding cyst until only a small number of basal stalk cells remained at its posterior (Fig. 11B and Movie S5). During this almost 15h viewing period, the number of cells posterior to the cyst declined from 32 to 13 (Fig. 11C). Fixed images of later stages, after two egg chambers have budded (84h APF; Fig. 11D) revealed an average of 9 cells in the stalk posterior to the terminal egg chamber, resembling the structure seen in newly-eclosed adults (Fig. 3A). Prior to budding, there were 50-65 cells posterior to the germline throughout the time range of 24 to 48h APF (Fig. 11A).

**Figure 11.**
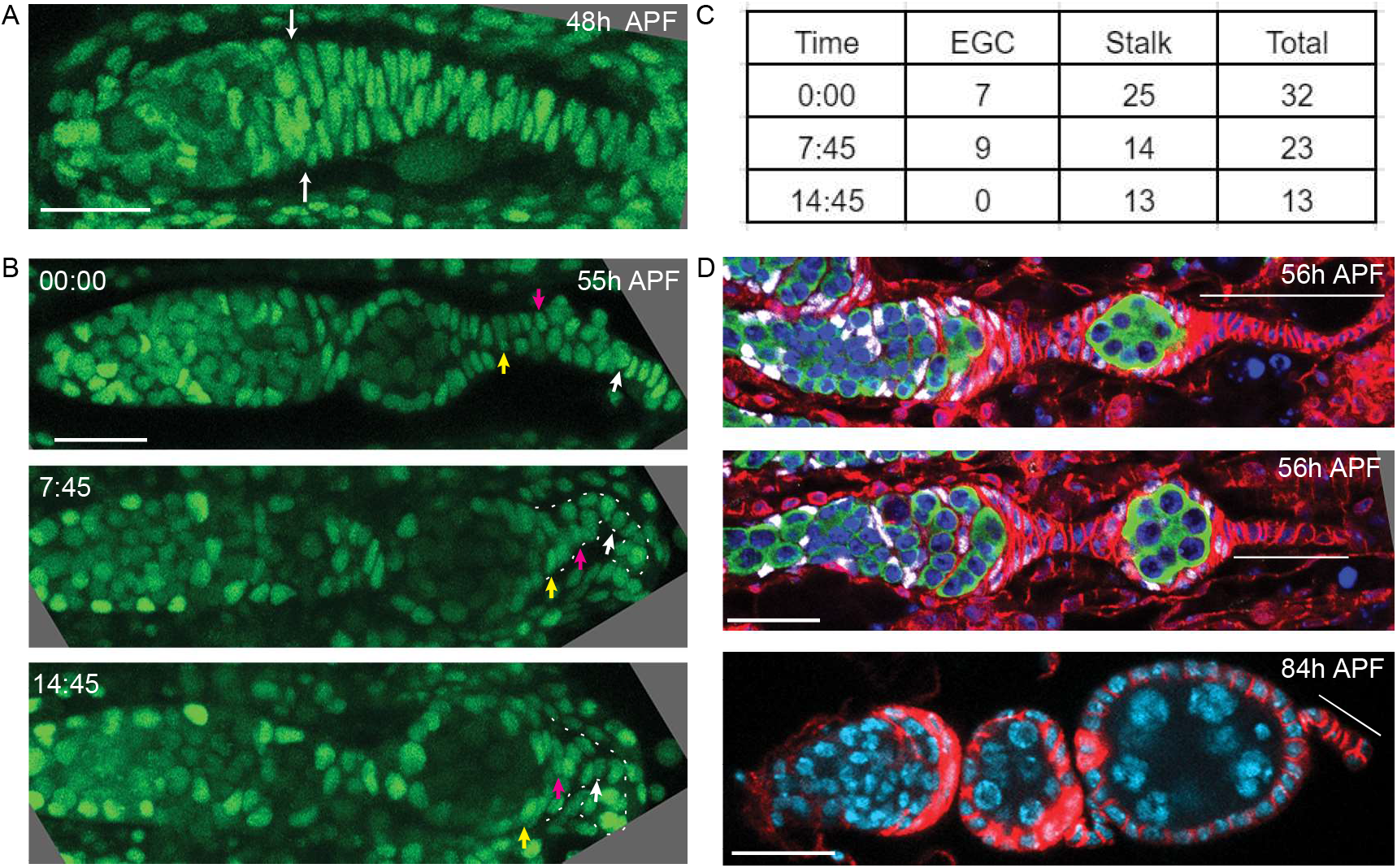
Basal stalk and EGC cells are the source of the first FCs. (A) Z-projection of a Ubi-GFP labeled germarium at 48h APF from live imaging to show the cells posterior to the germarium. White arrows indicate the posterior edge of the germline. (B) Stills from a movie of a 55h-APF germarium (Movie S6) show that the stalk, outlined by the white dotted line in the lower panels, progressively shortens as the first cyst moves posterior. Three cells were tracked: the white cell began in the stalk 13 cells away from the cyst and ended 4 cells away; the magenta cell was initially 7 cells away from the cyst and ended on the posterior of the cyst; and the yellow cell began as the first cell in the stalk and ended on the posterior third of the cyst. (C) Cell counts posterior to the germline for the germarium shown in (B). (D) Fixed images showing that the basal stalk becomes shorter over time. Thin white lines extend the length of the stalk. The top two images are stained for TJ (white), Vasa (green), DAPI (blue) and Fas3 (red). The lower image is stained for DAPI (blue) and Fas3 (red). After 2 egg chambers have budded there are an average of 9 cells in the stalk (n=12 stalks; the example in D has 9 cells). Scale bars, 20 µm.

Thus, it appears that many cells in the basal stalk at 48h APF end up as FCs on the first budded egg chamber, together with EGC derivatives. Some EGC cells likely pass over the entire budded cyst to occupy the space between the germarium and the budded cyst, and may be joined there by some of the cells that were the most posterior ICs at 48h APF to form the secondary EGC. It is very unlikely that any ICs at 48h APF become FCs on the most mature egg chamber of newly-eclosed adults.

Imaging from about 60h APF onwards revealed the most posterior cyst in the germarium rounding up and moving a short distance into the secondary EGC, which includes dividing cells (Fig. S6A-C). Tracked cells on the first budded egg chamber moved short distances anterior as the stalk elongated between the first and second egg chambers (Fig. S6C). ICs anterior to the second budding cyst moved posterior as the germarium elongated and appeared poised to associate with later cysts (Fig. S6A). We infer that the second budded egg chamber has FCs derived from the secondary EGC, which is itself derived from primary EGC cells and, likely, some posterior ICs at 48h APF. FCs on later egg chambers likely derive solely from cells that were ICs at 48h APF.

### Basal cell precursor contributions to the first-formed FCs and a basal stalk

Live imaging indicated that many cells in the basal stalk at 48h APF later become FCs on the first budded egg chamber, while a small number do not migrate over the cyst and form a short basal stalk structure of 5-10 single-file cells posterior to egg chamber 4 in newly-eclosed adults (Fig. 3A). Consistent with this scenario, when we examined lineages marked at 36h APF, we found 13 ovarioles with label in the basal stalk and in FCs of egg chamber 4 (Fig. 7F, G), suggesting a common origin from basal stalk cells at 36h APF. There were 18 ovarioles where FCs in egg chamber 4 were marked but the adult basal stalk was not, consistent with an origin from EGC or anterior basal stalk cells at 36h APF (Fig. 7E, H-J). The proportion of ovarioles containing marked FCs in egg chamber 4 that also contained marked FCs in egg chamber 3 was higher when marked basal stalk cells were absent (7/18) than when they were present (2/13), consistent with a more anterior origin of lineages that did not include the adult basal stalk (Fig. 7E).

In summary, live imaging together with lineage analysis initiated at 36h APF revealed a separate group of FC-only precursors, now identified as basal stalk and EGC cells posterior to the germarium, that become FCs of the first-formed egg chamber and contribute also to the second-formed egg chamber. Lineage analysis suggested there are additional FC-only precursors that contribute to more anterior FCs. Those precursors may be located partly in anterior regions of the EGC at 36h APF but it seems likely that most derive from ICs at 36h APF, given the observation that the germarium has an adult morphology after two egg chambers have budded, with no EGC structure. Although we could only track a subset of cells during live imaging, we did not find any evidence for cells posterior to ICs entering the germarium at any stage, consistent with all ICs present in the germarium after two cycles of budding originating from ICs at 36h APF. Importantly, the precursors of FSCs, which lie anterior to FC-only precursors, must therefore arise from cells within the germarium (ICs) throughout pupal development, whether they give rise to just FSCs and FCs or also to ECs.

### Pattern of precursor divisions along the AP axis

The analysis of MARCM clones in 2d-old adults provided information about the pattern of precursor divisions over time, based on the frequency of labeling specific precursors and the yield of adult cells from each precursor (Table 1). There was a severe reduction in EC-only precursor division rates from −4d to −2d (24h to 72h APF) with the frequency of such clones falling from 58% and 57% (−5d and −4d) to 30% (−3d) and then 14% (−2d). Moreover, the decline in division spread from the anterior; the yield of r1 ECs relative to r2a ECs declined from 2.2 (−5d) and 2.6 (−4d) to 0.9 (−3d) and 0.8 (−2d). The yield of marked cells per EC/FSC or FSC precursor was consistently higher than for EC-only precursors at all times of clone induction, indicating higher division rates in more posterior locations. These yields underestimate EC/FSC and especially FSC precursor output because they do not include FCs.

We also examined precursor cell cycling using EdU and FUCCI labeling. We incubated dissected pupal ovaries at various developmental stages with EdU *in vitro* over a 1h time period to detect cells in S phase. From 0-24h APF, ICs throughout the germarium often incorporated EdU (Fig. S1A-C). By 48h APF, the prevalence of anterior EdU-labeled cells was quite low, whereas EGC cells had frequent EdU signals (Fig. S1D). Even at 60h APF, however, an occasional IC in the anterior half of the germarium could be found with EdU label; no anterior ICs labeled by EdU were evident at 96h APF (Fig. S1E, F). At 96h APF some cells near the middle of the germarium but anterior to strong Fas3 expression, which may be r2a EC or EC/FSC precursors, did still incorporate EdU.

In the FLY-FUCCI system, RFP- and GFP-tagged proteins accumulate during the cell cycle under the influence of a chosen promoter, while degrons from CycB and E2F1 promote rapid degradation of RFP and GFP fusion proteins at the start of G1 and S phase, respectively [71]. While RFP and GFP are expected to be reliably absent during G1 and S phase, respectively, the speed of recovery of each signal during the next phase of the cell cycle depends on the strength of transcription of FLY-FUCCI components. We used *UAS*-driven *FLY-FUCCI* transgenes and found *C587-GAL4* to be more effective than *tj-GAL4* or *act-GAL4* in producing GFP and RFP FUCCI signals in the developing somatic IC population and in adult ovaries. In adult germaria, all r1 ECs express only GFP (Fig. 12G), indicating G1 (or a G0 state). r2a ECs include a mix of GFP-only G1 cells and cells with both GFP and RFP, indicating G2 (Fig. 12G). FSCs additionally include some cells with RFP-only (late S-phase) and no GFP or RFP (early S-phase, verified by co-labeling with EdU) (D. Melamed pers. comm.).

**Figure 12.**
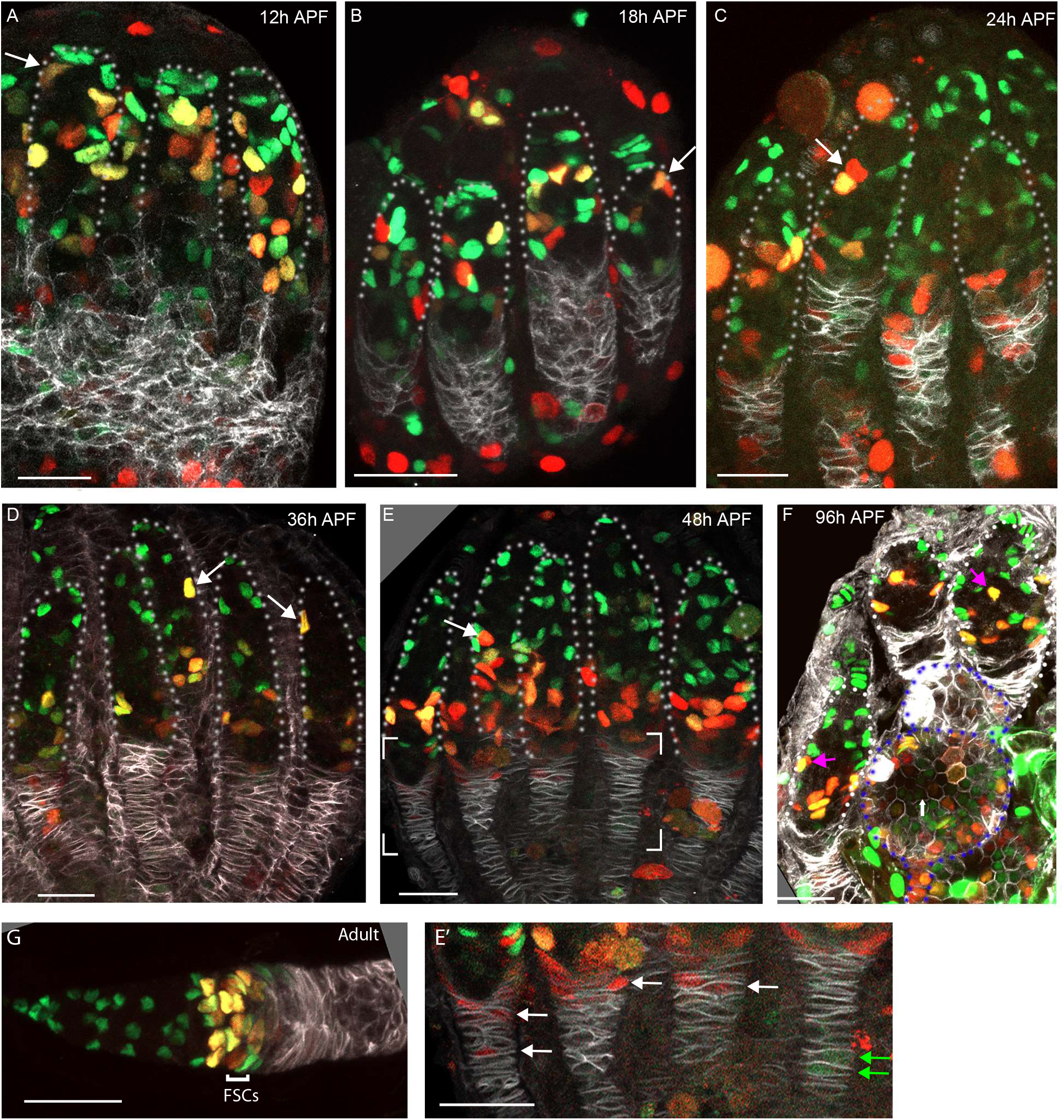
FUCCI labeling shows progressive accumulation of anterior ICs in G1 or a G0 arrest. (A-G) C587-GAL4 was used to drive UAS-FUCCI expression and ovaries were stained for Fas3 (white) and TJ (not shown) to identify all somatic cells. Somatic cells were either green (G1 or G0), red (late S phase), red/green (G2) or without any color (early S-phase for cells with adequate C587-GAL4 expression). Germaria are outlined with dotted lines. All bars, 20 µm. (A-F) Many red or red/green cells are present in anterior locations close to Cap Cells at (A)12h APF and (B)18h APF (arrows) but (C) by 24h APF and (D) 36h APF fewer such cells are evident and are further from the anterior (arrows). (E) At 48h APF ICs in the anterior third of the germarium are uniformly green (G1 or G0). The bracketed region is magnified with enhanced colors to reveal (E’) that many cells in the EGC are red (S-phase), while one or two more posterior, basal stalk cells are faintly green but C587-GAL4 expression is low at these locations. (F) At 96h APF, after three egg chambers have budded (two are outlined with blue dotted lines), red/green ICs can still occasionally be seen in the anterior half of the germarium (pink arrows) but (G) in adults this territory, occupied by r1 ECs, is entirely green, followed by a mix of green and red/green more posterior r2a ECs, and FSCs, which are predominantly red/green (G2), red or colorless (S-phase) and with only a few green (G1) cells. In adults, C587-GAL4 expression extends through all ECs and FSCs but ends near the FSC/FC border.

From 12-36h APF, ICs at all AP locations in the germarium included some cells with each FUCCI color combination, although G1 cells were always most frequent in more anterior regions (Fig. 12A-D). By 48h APF, almost all somatic cells in the anterior half of the germarium were in G1 or G0 (GFP-only), consistent with substantially reduced cycling (Fig. 12E). A large fraction of EGC cells were in S phase (marked by RFP only) (Fig. 12E’). By 96h APF, after two or three egg chambers have budded, the *C587*-expressing domain had retracted anteriorly, roughly to territory anterior to strong Fas3 staining and the pattern of FUCCI labeling was quite similar to adults, with anterior cells uniformly in a G1 or G0 state, followed posteriorly by cells largely in G1 and G2 and then some cells in S phase also (Fig. 12F). Thus, direct observation of EdU incorporation and cell cycle markers confirms the progressive decline in anterior IC proliferation over time that was inferred from lineage analyses and suggests that the anterior to posterior gradient of proliferation stretches all the way from the most anterior ICs to posterior ICs and EGC cells, which appear to cycle rapidly.

### A putative FSC/FC precursor marker yields lineages that also produce ECs

The location of precursors of a specific cell type can sometimes be deduced by taking advantage of restricted gene expression patterns to target limited sets of precursors for lineage marking. However, such approaches commonly suffer from limited targeting specificity and often do not allow single-cell lineage tracing [72, 73]. If cell marking is not uniformly precise, it is challenging to link a spectrum of initially marked cells with a spectrum of lineage outcomes because the same sample cannot be examined at both the start and the end of an experiment to infer precursor-product relationships directly. Recent single cell RNA sequencing of developing ovaries in late third instar larvae led to the discovery of a variety of genes with regional expression patterns, providing resources for targeted cell labeling [51]. The authors identified a cell population posterior to ICs that preferentially expressed *bond* mRNA prior to pupariation. A *bond-GAL4* driver was then used together with a temperature-sensitive *GAL80* transgene to initiate GFP-marked lineages using standard G-trace reagents [74], keeping animals at 29C during larval stages to initiate labeling, switching to 18C (to terminate labeling) about one day after pupariation and scoring marked cells in 2d-old adults [51]. Most of the resulting labeled ovarioles (31/32) were described as including marked FSCs or FCs. FSCs were not scored directly but were inferred from the presence of marked FCs in the germarium and youngest egg chambers of 2d-old adults, and the fraction of 31 samples specifically with marked FSCs, rather than just marked FCs, was not specified. More than half of the labeled ovarioles also contained marked ECs (19/32). *bond-GAL4* expression was noted to be less specific than *bond* RNA, with some expression in ICs in late 3^rd^ instar larvae, while *bond* RNA was seen expressed throughout the IC region in earlier 3^rd^ instar larvae. Although the majority of ovarioles with marked FSCs evidently also contained marked ECs, the authors suggested that separate precursors in these developing ovarioles gave rise to ECs and to FSCs, with the latter situated posterior to ICs in late third instar larvae. Since both of these conclusions are directly contradicted by our single-cell lineage analyses and morphological studies, we investigated lineages targeted by *bond-GAL4* using reagents donated by the authors. We scored results in newly-eclosed adults rather than 2d-old adults in order to count FSCs directly, rather than inferring their presence indirectly, and to include also all FCs that are produced from precursors during pupation.

We found a high frequency of GFP-marked cells when animals were kept at 18C throughout (37 labeled ovarioles out of 76), suggesting “leaky” clone initiation, with marked ECs or ECs together with FSCs present in almost all cases (35/37) (Fig. 13A-D). We saw no increase in the proportion of ovarioles with marked ECs or FSCs when larvae were raised at 29C before being moved to 18C about 2d after pupariation (eclosing 6d later; 13 of 44 ovarioles labeled), or when animals were raised at 18C before shifting to 29C for the final 3d or 4d of pupation (23 of 88 ovarioles labeled) (Fig. 13D). Because we observed no marked temperature dependence for these EC/FSC-containing clones we cannot define the time at which cells were first marked. However, we expect that most lineages were initiated close to the larval/pupal transition based on the composition of the lineages (see below) and because TF and Cap cells were occasionally labeled.

**Figure 13.**
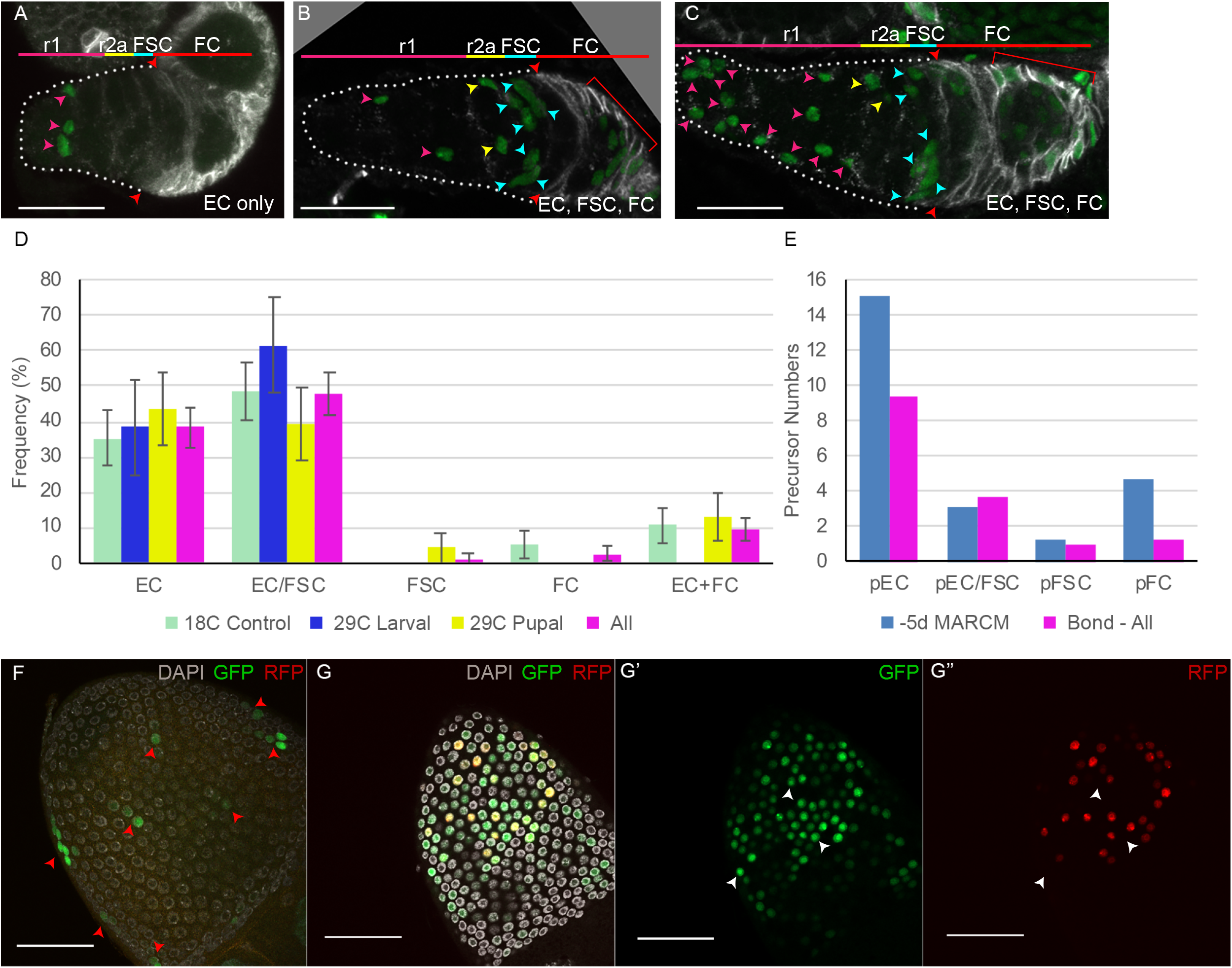
Lineages induced by bond-GAL4 reveal typical IC precursor patterns and a later EGC contribution. (A-G) bond-GAL4/G-TRACE clones (green) were examined in 0-12h adult ovaries, stained with antibodies to Fas3 (white in (A-C)) and DAPI (white in (F, G)). (A-C) For animals maintained at 18C throughout development, we observed mainly (A) marked ECs only or (B, C) marked ECs, FSCs and FCs; scale bars, 20 μm. (D) Distribution of clone types observed for animals kept at 18C throughout (green), raised at 29C and moved to 18C 2d after pupariation (“29C larval”, blue), raised at 18C and moved to 29C for the last 3-4d of pupation (“29C pupal”, yellow), or the aggregate of all results (pink). EC/FSC and FSC categories almost always included FCs also. (E) Estimates of precursor numbers for MARCM clones induced 5d before eclosion (blue) and for the aggregate of all bond-Gal4 lineages (pink) after disaggregating raw data into single lineages (see Methods). (F) Posterior egg chamber (29C larval) with small GFP clones (arrowheads). Multiple z-sections are projected to show all clones. (G) Posterior egg chamber (29C pupal) with patches of (G’) GFP-positive cells, some of which also express (G”) RFP, indicating relatively recent expression of bond-GAL4. (F-G) Scale bars, 50 μm.

The marked lineages were similar in their composition of cell types (Fig. 13A-D) to those we described earlier using *hs-flp* to initiate clones at pupariation (5d before eclosion), suggesting that they may have been induced at a similar time. The majority of ovarioles from the *bond-GAL4* experiments (considering all three temperature conditions together) included either only marked ECs (about 40%) or marked ECs and FSCs (about 50%, almost all together with marked FCs) (Fig. 11D). Only one out of 73 labeled ovarioles had marked FSCs (and FCs) but no marked ECs. These results clearly provide no direct evidence of the labeling by *bond-GAL4* of precursors that produce FSCs and FCs without also producing ECs.

The number of labeled ECs and FSCs (when present) per germarium was consistently higher than we observed with *hs-flp* induced MARCM or multicolor clones induced at pupariation (Fig. 13A-C; Table 3), suggesting that most labeled ovarioles in the *bond-GAL4* experiment contain lineages derived from more than one precursor. The number of marked cells per ovariole was roughly consistent with the output from two single-cell lineages in MARCM clones initiated at 0h APF. We therefore calculated the component single clone types (frequency and number of products) that would produce the observed *bond-GAL4 > G-trace* results if every ovariole had exactly two marked lineages (see Methods). The inferred single precursor lineage types were similar to those induced by *hs-flp* at 0h APF (Fig. 11E; Table 3). The most significant differences were a greater yield of ECs per EC-only lineage and fewer FC-only lineages for lineages induced by *bond-GAL4*; both differences are consistent with slightly earlier clone initiation. It is therefore likely that *bond-GAL4* labeled precursors sporadically throughout the IC region, mostly shortly before pupariation. In keeping with our other lineage studies, inferred single cell lineages that contained marked FSCs, also included marked ECs in almost all cases. The source of lineages reported to include FSCs in the earlier study [51] may also have been pre-pupal ICs. Moreover, the ovarioles with marked ECs and FSCs may have derived from common precursors, consistent with all of our studies, rather than a combination of EC-specific and FSC-specific precursors, as suggested by the authors [51].

When animals with *bond-GAL4* and *G-trace* were maintained at 29C during the last 3d or 4d of pupation we observed an additional type of lineage, either in isolation or together with the types of marked cells described above (55/88 ovarioles; 0/108 when at 18C throughout) (Fig. 13G). The second lineage pattern consisted of a few isolated FCs or small FC clusters in the most mature egg chamber, suggesting several independent recently initiated lineages. Some of the GFP-labeled FCs also expressed *UAS-RFP*, indicating current or recent *bond-GAL4* activity (Fig. 13G). These patterns were not reported previously [51] because the first-formed egg chambers are no longer present in 2d-old adults. Surprisingly, the same type of clone was observed if animals were shifted from 29C to 18C about 2d after pupariation (30/44 ovarioles, with only 6 ovarioles also including RFP-expressing FCs) (Fig. 13F). The individual patches of marked FCs were much smaller than observed for lineages induced by *hs-flp* at 36h APF and therefore presumably result from induction of multiple lineages at a later time due to perduring *bond-GAL4* or *UAS-flp* products. The second pattern of lineages initiated by *bond-GAL4* suggests selective expression of that marker in FCs of the first-formed egg chamber in late pupae and adults. We have no clear evidence to evaluate whether *bond-GAL4* might also be expressed sufficiently in precursors of those FCs to initiate lineages earlier in pupal development, reflecting the posterior domain of *bond* RNA expression observed in late third instar larvae [51].

## Discussion

An important general objective, exemplified here by FSCs, is to ascertain the origin of adult stem cells and surrounding niche cells in order to understand how their number and organization are instructed during development. Our investigations of changing morphology, cell movements and cell lineage reveal a gradual and flexible adoption of these cell types according to AP location throughout pupal development, rather than a series of discrete, definitive cell fate decisions (Fig. 6). This contrasts with the development of many other types of differentiated adult cells.

Specifically, adult EC niche cells and FSCs develop from a common group of dividing precursors (ICs) that are interspersed with developing germline cysts. ICs express TJ and are located posterior to Cap Cells, which serve as adult niche cells for both GSCs and FSCs, and are already specified by the start of pupation. The most anterior ICs become regional subsets of ECs (r1 and r2a), which likely have distinctive roles in the adult [75, 76]. These precursors markedly attenuate division, starting from the most anterior locations, during the second half of pupation (Table 2, Fig. 12 and Fig S1). ICs further from the anterior produce both ECs (mainly r2a) and FSCs, but EC production declines over time (Fig. 7). More posterior ICs produce FSCs and FCs (Fig. 6), while the most posterior ICs produce only FCs . The deduced pattern of germarium development ensures the juxtaposition of ECs and FSCs. The use of common precursors likely contributes to producing those cell types in appropriate proportions, while EC numbers are limited by a progressive decline in proliferation that spreads from the anterior.

Germarium development is likely driven in part by the early establishment of TF and Cap Cells as a source of organizing signals but the behavior of adult FSCs, including conversion to FCs, also relies on a signaling source posterior to the germarium [24, 26]. A key signal comes from specialized FCs, known as polar cells, on the most recently budded egg chamber (Fig. 1J). At the start of pupation there are no budded egg chambers, so formation of the first FCs and the first egg chamber must proceed by a different route. We found that those first FCs derive from precursors that are posterior to ICs and differ from ICs during the first 48h of pupation by expressing Fas3 strongly, by lack of association with germline cells and by being restricted to produce only FCs, and not ECs or FSCs. Thus, the first population of posterior niche cells, and the most posterior portion of the ovariole of a newly-eclosed adult are exceptional in being derived from precursors that do not overlap with the precursors of adult FSCs.

Overall, almost the entire somatic cellular complement of the adult tissue that is subsequently maintained by stem cell activity, derives from precursors that also produce adult FSCs. Preliminary understanding of other adult stem cells suggests this arrangement may be common but not universal. There is evidence that both adult mouse gut stem cells and adult neural stem cells of the dentate gyrus derive from a common precursor pool that also gives rise to the adult epithelial tissue maintained by the stem cell [77, 78]. By contrast, subventricular zone neural stem cells appear to be specified during embryogenesis and remain quiescent, while the adult tissue it can replenish is built from other precursors [79, 80].

### Defining and locating EC, FSC and FC precursors through pupal development

This study is the first to examine Drosophila ovary development comprehensively during pupal stages through lineage analyses initiated at different times, and by systematic analysis of fixed specimens and live imaging. Each experimental approach included some challenges and leaves some issues unresolved but several key findings appear conclusive and supported by all three approaches.

The challenge of labeling only a single precursor among many is common in lineage studies. Here it was met by using very mild heat-shocks to initiate clones through *hs-flp* induction and a multicolor labeling technique that reduces the frequency of a specific color combination substantially relative to single-color clones. We were also able to evaluate the frequency of single-cell lineages from both the overall frequency of ovariole labeling and the frequency of clone patterns (exceptional, discontinuous clones), allowing us to derive mathematical estimations of strictly single-cell lineage frequencies and yields. The results led to robust conclusions that many precursors at the start of pupation yield only ECs, a smaller proportion yield both ECs and FSCs (as well as FCs) and very few produce FSCs with no ECs. Common precursors for ECs and FSCs persist for at least the first 72h of pupation (later times were not tested), although they are exceeded in number at 72h APF by precursors that produce FSCs but not ECs. All such FSC lineages also included marked FCs in newly-eclosed adults.

By 72-84h APF, after two egg chambers have been produced, the organization of the germarium, including the pattern of Fas3 staining, appears similar to the adult germarium. It is therefore plausible that the behavior of FSC precursors during the remainder of pupation is regulated just as in adults, reflecting, in effect, completion of germarium development up to two days before eclosion. The activity of common precursors for ECs and FSCs during the first half of pupation reflects a very different process in a different environment, with progressive germline cyst differentiation being organized simultaneously and no budded egg chambers to provide a key posterior signaling center. The existence of common precursors for FSCs and ECs during the first half of pupation could not therefore have simply been inferred or predicted on the basis of adult FSC behavior. Indeed, this finding, based on painstaking lineage analyses, contradicts a previous conclusion [51]. The prior conclusion was based on an approach that we also investigated, where marked lineages generally initiated from more than one precursor in an ovariole, and generally also produced marked ECs when marked FSCs were present. These results were previously interpreted as reflecting separate EC and FSC lineages in the same ovariole in accord with a presumption of early fate specification based on differential gene expression [51], but, in our hands, fit quantitatively with an origin from common EC/FSC precursors at the start of pupation, in accord with all of our other lineage studies.

Our lineage studies provided some information about precursor locations. Each lineage was restricted in its AP extent, but the whole collection of lineages included overlapping domains that spanned the whole length of the ovariole of a newly-eclosed adult, from the most anterior r1 EC to FCs and the basal stalk of the terminal egg chamber. Those observations strongly suggest that precursors are arranged in a continuum along the AP axis throughout pupal development (Fig. 6). The anterior and posterior limits of the EC/FSC/FC precursor domain were deduced from direct observation of pupal development. Clone marking by heat-shock induction of Flp recombinase is through mitotic recombination and can occur in any dividing cell. We directly observed EdU labeling in the most anterior ICs in early pupae, so we can deduce that the most anterior of the inferred continuum of precursors lie immediately adjacent to cap cells and give rise to adult r1 ECs. The posterior extent of these precursors was deduced by direct observation, including live imaging, of the production of the first FCs, as summarized below.

### Live imaging reveals the posterior origin, outside the developing germarium, of the first FCs

The precursors that give rise to FCs surrounding the most posterior egg chamber of newly-eclosed adults were identified by a combination of fixed and live images. These cells all express Fas3 strongly by 24h APF and comprise two distinguishable groups according to their locations and TJ expression prior to egg chamber budding. We named one group as Extra-Germarial Crown (EGC) cells in recognition of their location and appearance. EGC cells accumulate from a handful to over twenty during the first 48h APF. The second group of cells are further posterior, do not express TJ and are largely intercalated into a stalk of 1-2 cell’s width at 48h APF prior to egg chamber budding. During budding, the most mature germline cyst leaves the germarium largely devoid of accompanying ICs, which do not yet express Fas3 strongly. The cyst then enters EGC territory and moves a considerable distance from the germarium as it is enveloped by EGC and basal stalk cells.

Towards the end of the budding process, cells posterior to the new egg chamber are stacked in single file, forming a basal stalk, which is stably retained into adulthood, confirming and supplementing prior descriptions of basal stalk formation [45]. We observed several lineages that included labeled cells in the adult basal stalk and in a FC patch on the terminal egg chamber, reflecting contributions of pupal basal stalk cells. Meanwhile, some EGC derivatives also remain on the first budded egg chamber, while others migrate beyond the budded cyst to form a secondary EGC between the germarium and the budded egg chamber. Secondary EGCs provide cells to partially coat the second cyst to emerge from the germarium. The second cyst appears from live imaging to move out of the germarium together with some associated ICs. Lineage studies also suggest that FCs of the second budded egg chamber derive from both EGC cells (producing FCs in egg chamber 4 and 3) and ICs (producing FCs in egg chamber 3 and more anterior locations) (Fig. 7E). The budding of the third, and subsequent, egg chambers is likely similar to the process in adults, with cysts emerging from the germarium completely encased by FCs, as there is no longer a recognizable EGC after two egg chambers have budded and the two most mature germline cysts are associated with strongly Fas3-positive cells within the germarium after the second egg chamber is about to bud.

These observations and deductions not only reveal the posterior limit of EC, FSC and FC precursors as including basal stalk cells at about 48h APF, but also provide evidence that FSCs derive from precursors that are ICs. The evidence is temporal and numerical. First, we saw an expansion of the EGC population from 12-48h APF prior to egg chamber budding (Fig. 8). During that time and during the budding process, we did not observe any cells from the EGC moving into the germarium. Moreover, as soon as the second egg chamber is about to bud, it is highly likely that the FCs of both of the next two future egg chambers are the Fas3-positive IC cells that surround those cells in the germarium. Those prospective FCs therefore almost certainly derive from precursors that were never outside the germarium. FSCs derive from precursors that are further anterior, so those precursors were even more certainly always within the germarium as ICs. Second, just as the first germline cyst leaves the germarium at, or shortly after 48h APF, there are over 60 TJ-positive somatic ICs in the germarium (Fig. 8D). At this time, many EC/FSC precursors are still dividing and not all future ECs have terminated divisions. The total number of ECs (about 40 on average) plus FSCs (about 16) in an adult germarium is less than 60, so it is clear that the ICs in the germarium at this time must include all EC and FSC precursors and a significant number of FC precursors.

### Deductions from targeted lineage analyses

A recent study used single cell RNA sequencing to explore the diversity of precursors in late larval ovaries. Of most relevance here, it was suggested that a distinct group of cells posterior to ICs and characterized by *bond* expression give rise to FSCs and FCs but not ECs [51]. As described earlier, the supporting lineage studies and our reproduction of lineages initiated from cells expressing *bond-GAL4* cannot be interpreted definitively because ovarioles almost never contained lineages derived from a single cell. The results are nevertheless compatible with all of our other observations demonstrating that FSC precursors at the start of pupation also give rise to ECs, and that FSC precursors are ICs, residing within the developing germarium throughout pupation. Indeed, because we scored FSCs directly, counted the number of labeled ECs and FSCs in each sample, and could compare the results to those we had already obtained for single-cell lineages initiated at pupariation, we were able to deduce that ovarioles almost always harbored more than one lineage initiated by *bond-GAL4*. This allowed us to estimate the composition of single-cell lineages and found they were very similar to those induced by heat-shock induction of marking in all dividing cells. Although temperature control of initiation of clones with *bond-GAL4* was leaky, we can infer that the FSC-yielding lineages mostly reflect initiation by *bond-GAL4* expression in ICs shortly before pupariation. This is compatible with the observation that *bond-GAL4* is expressed at low levels throughout the IC region in mid-third instar larvae [51].

We also observed another type of lineage when *bond-GAL4* activity was increased by raising the temperature during pupal development. Here, labeled FCs were seen in small patches on the first budded egg chamber. Those lineages would not have been apparent in the earlier study because only 2d-old adults were examined [51]. The labeled FC patches were much smaller than those induced by lineages initiated at 36h APF, so they must reflect *bond-GAL4* expression in those FC precursors late in pupation. We did observe sporadic *bond-GAL4* expression in FCs of the most mature egg chamber in adults. It seems plausible that the expression of *bond* RNA observed posterior to ICs in late larvae [51] may continue in an analogous location, posterior to the germarium, in the EGC or basal stalk, and as those cells become FCs. However, at present, we have no direct evidence of *bond* expression during pupation or of lineages initiated in cells posterior to ICs prior to budding of the first egg chamber.

### Coordination between germline and somatic ovarian cell development

Ovary development not only organizes suitable numbers of FSCs, GSCs and their niche cells into an adult germarium, ready to fuel lifelong adult oogenesis, but also initiates oogenesis very early, so that the first eggs can be laid shortly after eclosion. How is the posterior progression of increasingly mature cysts, starting from single anterior cells, organized?

Ecdysone acts positively in somatic cells of late third instar larvae to initiate GSC differentiation, manifest by induction of the differentiation factor, Bag-of-marbles (Bam) and then cystocyte formation, revealed by the transition of the rounded spectrosome to a branched fusome [59, 60]. Prior studies did not reveal whether Bam-GFP expression or PGC division initiates simultaneously in all PGCs distant from Cap Cells or in an anterior-posterior gradient, or how germline differentiation proceeds thereafter. We found that there is a clear anterior to posterior polarity of increasingly developed cysts that produced a mature steady-state pattern by 48h APF. This was deduced by counting the number of germline cells linked by fusomes, with 16-cell cysts first appearing 18h APF, and by observing the exclusion of EdU DNA replication signals from the most posterior regions of the germarium from 36h onwards, indicating occupancy by only 16-cell cysts. Moreover, a single posterior sixteen-cell cyst emerged for the first time between 36h and 48h APF.

These observations raise the question of what underlies the spatially organized initiation or progression of differentiation of PGCs that are in different locations at the larval/pupal transition. The BMP signals that maintain anterior pre-GSCs are likely highly spatially restricted, as in adults [81], while hormonal Ecdysone signals may be spatially uniform, suggesting that there must be other important differentiation signals. Potential sources of relevant signals are the adjacent somatic cell precursors, which will later become ECs. There is some evidence that the ordered posterior movement of germline cysts in adult germaria depends on EC interactions [82]. During pupal development, the situation is different because the initiation of germline differentiation must be spatially organized and many EC precursors are still dividing during this time. It will be interesting to investigate whether pupal EC precursors or adult ECs have spatially graded adhesive properties or send graded differentiation signals according to their AP position.

### Organizing patterned development of adult stem cells and niche cells

It is common to think of developmental processes and cell types in terms of establishment of transcriptional networks and key markers that bear witness to permanent decisions. Investigations of adult FSCs, and other adult stem cell paradigms such as the mammalian gut, as well as the observations described here regarding FSC development, reveal a more flexible process, in which environmental signals reflecting current cell locations may be more significant than stable expression of master regulators of transcription. For example, the transcription factor TJ is selectively expressed in many developing and mature ovarian somatic cells, and it has been shown to be important for the specification of Cap Cells and many IC derivatives [52, 54]. We therefore suspected that EGC cells, posterior to ICs, might be FC precursors because they also expressed TJ. Although that suspicion was supported by live imaging studies, the same studies showed that TJ-negative basal stalk cells also became FCs. Thus, TJ neither defines ICs nor EC/FSC/FC precursors. We think it is likely that studying the spatial distribution of external signals and their interpretation by somatic cells in the developing ovariole will be essential to understand why some cells initially mix with germline cells to become ICs, while others do not, and how these cells become ECs, FSCs and FCs just in time to serve the appropriate functional role.

The signals that guide formation of Fas3-positive FCs in the adult germarium include low Wnt pathway activity and high JAK-STAT pathway activity. The location of FC differentiation is set, at least in part, by a Wnt pathway activity gradient that declines from the anterior and by the opposite polarity of graded JAK-STAT pathway activity initiated by a ligand derived from polar cells of the previously budded egg chamber [24, 26]. Prior to budding of any egg chambers in pupal ovaries, there is no polar FC source of JAK-STAT pathway ligand. It will be interesting to discover whether differentiation of the first FCs in pupal ovaries also relies upon being distant from an anterior source of Wnt and perhaps other pathway ligands, such as Hedgehog, and whether a JAK-STAT signal, coming from a source other than polar cells, is employed.

## Supporting information

Supplemental Figures and Description of Movies

## Acknowledgments

This work was supported by NIH RO1 GM079351 to DK. We thank Ruth Lehmann, Dorothea Godt, Erika Bach, Ramanuj DasGupta and Ward Odenwald for reagents, Pegah Kohosravi-Kamrani, Shay Mallick, Aaron Choi and Jack Misner for help with experiments and analysis, David Melamed for continued discussions and input, the Bloomington stock center for provision of genetic reagents, the Developmental Studies Hybridoma Bank (DSHB) for antibodies, FlyBase as an information resource, and the confocal microscope resource provided by the Dept. of Biological Sciences, Columbia University.

Conceptualization, A.R. and D.K., Formal Analysis, D.K., Funding Acquisition, D.K., Investigation, A.R., H.K., R.M. K.P., Methodology, H.K., K.P., A.R., R.M. D.K., Supervision, A.R. and D.K., Validation, A.R., H.K., R.M., K.P., D.K., Visualization, A.R., H.K., R.M., Writing-Original Draft, D.K., Writing-Review & Editing, A.R., H.K., R.M., K.P., D.K.

## Materials and Methods

### Staging of pupae

Third-instar larvae were sorted to select females, transferred to a vial with food and checked every hour to mark the time each individual developed into a puparium (white, but immobile, with small anterior spiracles). This time is 0 hours APF (after puparium formation). We found that keeping animals in the light at night and in the dark during the day on a 12h/12h cycle, allowed more of them to pupate during the daytime. We used a programmable outlet timer (Nearpow) together with a manual LED soft white nightlight (Energizer cat. 37099) installed in a dark incubator at 25C.

### Dissection and staining of pupal ovaries

Pupae were removed from the vial wall by adding a drop of water and transferred with forceps into a well of a Corning PYREX glass spot plate (Corning, cat. no. 7220–85) containing either PBS or 4% paraformaldehyde solution with care taken not to damage or pierce the posterior end of the pupa. If the dissection was done in fixative solution, it could take no longer than 10 min (ovaries were rocked in clean fixative for a minimum of 15 min, but for no longer than 30 min). Ovaries from 72h APF and older had fewer tears in them when dissected directly into fixative. Using a dissecting microscope, pupae were held against the bottom of the well using forceps and the posterior tip was cut off using Vannas Spring Scissors (2.5mm straight edge, Fine Science Tools, Foster City, CA, item number 15000-08) about a third of the way in from the posterior end. The anterior part of the pupa was removed to a discard well. The posterior end was searched for ovaries (clear, spherical, striated structures) by gently dislodging fat tissue surrounding the ovaries so as not to cloud the well with debris. This was done by gently tearing apart the contents of the pupal case or swirling them in the well using two pairs of forceps. As soon as they were spotted, ovaries were transferred to a new well of fixative by the following process. A 20μl pipette tip with a pipettor set to 20μl was coated in 10% Normal Goat Serum (NGS) by pipetting up and down several times. The pipettor was set to 5μl to pipette up the ovary from the dissection well to transfer into the fixative well. Ovaries were fixed by rocking in the well for 15 min at room temperature. Fixative was removed with a 1000μl pipette set to 270μl to slowly and carefully pipette up the liquid. The 1000 μl pipettor was used to remove liquid after all subsequent steps. Ovaries were rinsed 3x in 2% PBST 0.5% Tween solution for 5, 10, and 45 min at 4C by rocking very slowly in a horizontal plane. The glass dissection dish was covered with a large pipette tip box. Ovaries were rinsed for 30-60 min in 10% NGS solution at 4C, and then rocked gently in primary antibody overnight. Ovaries were rinsed 3x with 0.5% PBST and then incubated for 1 hour (covered, cold room, rocker set on 2) in secondary antibody diluted 1:1000 with 0.5% Triton. Ovaries were rinsed 2x with 0.5% PBST and 1x with PBS. To mount ovaries, a 20μl pipette tip coated in 10% NGS solution as described above was set to 5μl and used to capture the ovary and transfer to a glass slide. 20μl of DAPI Fluoromount was added to a coverslip, and then placed on top of the slide with the ovaries. Care was taken to avoid pressing the coverslip and potentially disfiguring the mounted ovaries. For older pupal ovaries, specifically 72h APF and onwards, forceps were used to gently tear apart the ovarioles to separate germaria for clearer imaging. A sharpie was used to draw arrows to mark the location of the ovaries on the coverslip.

### EdU labeling

Ovaries were dissected directly into 15μM EdU in Schneider’s insect medium, incubated for 1h at room temperature and then fixed for 10 min at room temperature in 4% paraformaldehyde. EdU incorporation was detected using the Click-iT Plus EdU Imaging Kit C10637 (Life Technologies).

### Immunohistochemistry

Monoclonal antibodies against LaminC, Fasciclin III, Vasa and Hts were obtained from the Developmental Studies Hybridoma Bank, created by the NICHD of the NIH and maintained at The University of Iowa, Department of Biology, Iowa City, Iowa 52242. 7G10 anti-Fasciclin III was deposited to the DSHB by C. Goodman, and was used at 1:300 for multicolor lineage experiments and at 1:250 in all other stainings. 1B1 anti-Hts was deposited by H.D. Lipshitz. LC28.26 Anti-LaminC at 1:50 was deposited by P.A. Fisher. Anti-Vasa (used at 1:10) was deposited by A. C. Spradling/D. Williams and used for staining adult ovaries. For pupal ovaries, the monoclonal rat anti-Vasa was not effective and instead rabbit anti-Vasa (gift from R. Lehmann, NYU medical school) was used. Other primary antibodies used were rabbit anti-Zfh-1 at 1:5000 (a gift from R. Lehmann, NYU medical school), guinea pig anti-Traffic Jam at 1:5000 (a gift from Dorothea Godt, University of Toronto, Canada), anti-β-galactosidase (Catalogue No. 55976, MP Biomedicals) at 1:1,000; anti-GFP (A6455, Molecular Probes) at 1:1,000; goat FITC-anti-GFP (Abcam ab6662) at 1:400. Secondary antibodies were Alexa-488, Alexa-546, Alexa-594 or Alexa-647 from Molecular Probes. Ovarioles from multicolor experiments were mounted in Prolong Gold Antifade (Invitrogen). DAPI-Fluoromount-G (Southern Biotech) was used as mounting medium for all other experiments. Images were collected with a Zeiss LSM700 or LSM800 laser scanning confocal microscope (Zeiss) using a 63x 1.4 N.A. lens.

### Adult ovary fixation and staining

Ovaries were fixed in 4% paraformaldehyde in PBS for 10 min at room temperature, rinsed three times in PBS with 0.1% Triton and 0.05% Tween-20 (PBST), and blocked in 10% normal goat serum (Jackson ImmunoResearch Laboratories) in PBST. Ovarioles were incubated in primary antibody for 45 min for multicolor lineage experiments and overnight for all other experiments. Ovarioles were rinsed in PBST three times and incubated for 1 h in secondary antibodies diluted 1:1000.

### Live imaging and analysis

Live imaging was performed as in [70], except that pupal ovaries dissected in imaging medium were left intact instead of separating into individual ovarioles. Intact pupal ovaries were mixed with Matrigel in a well of a gas-permeable imaging chamber. Ovaries were of genotype *yw hs-flp; ubi-GFP FRT40A* (Ubi-GFP ovaries) or *yw hs-flp/yw; ubi-GFP FRT40A FRT42B His2Av-RFP/tub-lacZ FRT40A FRT42B* (referred to as multicolor ovaries). Clones were generated in the multicolor ovaries by two 10 min heat shocks at 37C spaced 8 hours apart, 48 to 72 hours before imaging in order to generate GFP-only and RFP-only cells that were easier to track. Images were acquired every 15 min with 2.5 μm between Z-stacks. Volocity Software (Quorum Technologies, Puslinch, Ontario, Canada) was used to make the 4D reconstruction of the UbiGFP labeled ovary. All nineteen z slices covering 45 μm were used in the reconstruction; however, germaria drifted during imaging such that FCs covering approximately one quarter of the budded egg chamber on the lower germarium were not imaged. Cells were colored with Procreate software (Savage Interactive, North Hobart, Australia). Germline cells in the UbiGFP movie were identified based on their larger size and paler labeling with GFP compared to somatic cells.

### Clonal analysis and FUCCI labeling genotypes

MARCM clones for lineage analysis were generated on 2R and 3R using the following genotypes: (2R) yw *hs-Flp, UAS-nGFP, tub-GAL4/ yw; FRT42D act-GAL80 tub-GAL80 / FRT42D sha; act>CD2>GAL4/ +* and (3R) yw *hs-Flp, UAS-nGFP, tub-GAL4/ yw; act>CD2>GAL4 UAS-GFP / +; FRT82B tub-GAL80/FRT82B NM*

For multicolor clones flies were of the genotype: *yw hs-flp/yw; ubi-GFP FRT40A FRT42B His2Av-RFP/tub-lacZ FRT40A FRT42B*

Genotypes for FLY-FUCCI were *C587-GAL4/ yw; UAS-FUCCI/Cyo; UAS-FUCCI/ (UAS-FUCCI or TM6B)*.

### Lineage analysis by MARCM

Pupae were heat shocked at 33C for 12 min at the number of days indicated before eclosion. Flies were dissected on the day of eclosion (0d MARCM) or collected on the day of eclosion and dissected 2 days later. Ovaries were stained using antibodies to Vasa, Fas3 and GFP. Cells were scored as r1 or r2a based on measurements that were performed to determine the lengths of these regions. To determine the boundaries of each region we examined EdU incorporation in ovaries of newly eclosed flies and measured the distance from cap cells to the end of Region 1 as determined by the presence of cysts that incorporated EdU. Region 1 cysts are going through mitosis and Region 2a cysts have the complete complement of 16 germ cells. The length ratio of Region 1:2a is about 3:1 in newly-eclosed adults. To measure the percentage of each egg chamber covered by marked FCs, GFP was used to count the number of marked FCs and DAPI was used to count the total number of FCs in each z section.

### bondGal4/G trace experiments

*bond-GAL4/G-TRACE* flies (a gift of Ruth Lehmann, NYU medical center) were crossed at 18C and either kept at 18C, shifted after egg-laying to 29C and shifted back to 18C in mid-pupation (flies took 6 more days to eclose at 18C); or shifted to 29C 3 to 4 days before eclosion. Flies of genotype *UAS-RFP, UAS-flp, ubi>stop>GFP / +; bond-GAL4/ tub-GAL80(ts)* were dissected on the day of eclosion.

### Inferring clone types, FSC and EC numbers in newly-eclosed adults from scoring 2d-old adults

In order to make quantitative conclusions about the developmental process up to eclosion from scoring clonal outcomes in 2d-old adults we made the following assumptions and adjustments. We assumed that EC numbers and locations did not change during this 2d interval because ECs generally appear to be quite long-lived. ECs are, however, produced from FSCs during adulthood, so it is likely that a small number of marked ECs scored at +2d were produced from FSCs after eclosion. The proportion of marked ECs arising from FSCs is likely small based on known rates of FSC conversion to ECs [24, 40] and the observed high frequency of FSC clones with no ECs (44%) from −2d samples. Individual marked FSCs can be lost, mainly by becoming FCs, or amplified at high rates during adulthood [23]. If one or more marked FSCs were present at 0d but became an FC by +2d the resulting marked FCs would reside around the two germarial cysts or in the two youngest egg chambers because egg chambers bud roughly every 12h, allowing four cycles of FC recruitment over 2d. We therefore scored an ovariole as containing an FSC (classifying it as an FSC-only or FSC/EC lineage, according to EC content) if there were any marked FCs up to and including the second egg chamber, even if there were no FSCs at +2d. In those cases, we scored the number of marked FSCs as zero. On average, the number of marked FSCs should remain constant from 0d to +2d, so we should obtain a very good estimation of FSC numbers at 0d by scoring the numbers of marked FSCs at +2d, provided we include all examples where all marked FSCs were lost as containing zero FSCs. By using these guidelines, we could surmise the number of different types of clone at 0d and the number of constituent ECs and FSCs from data collected by scoring clones in 2d-old adults.

### Calculation of single-cell lineage frequencies assuming independent recombination events

The proportion of marked ovarioles with a lineage originating from a single marked cell is generally estimated by assuming that the probability of each dividing precursor to be labeled is independent. The estimate derives principally from the observed frequency of unlabeled ovarioles but it is also dependent to a small degree on the number of potential target cells. Different numbers of assumed target cells are used in the calculations below to demonstrate that estimated single cell frequencies are not greatly altered by this value.

For the −6d to −2d time course, the average frequency of labeled ovarioles was about 60%. If there were, for example, only 6 target cells and we assume a binomial distribution (independent probabilities) of labeling, the chance of any one cell giving a clone is p and the chance of 6 cells not giving a clone is (1-p)^6^. The observed no-clone fraction was 0.4, so (1-p)= 0.86 and p =0.14. Hence, the single clone frequency among 6 cells is 6p(1-p)^5^, and the frequency among labeled ovarioles is 1/0.6 fold higher (because 60% were labeled). Therefore, the expected frequency of single-cell lineages is 6×0.14x(0.86)^5^/0.6= 0.66.

If there were instead 15 target cells then p = 0.06 (because 0.94^15^= 0.4) and the frequency of single-cell lineages is 15×0.06x(0.94)^14^/0.6= 0.63.

If there were 30 target cells then p = 0.03 (because 0.97^30^= 0.4) and the frequency of single-cell lineages is 30×0.03x(0.97)^29^/0.6= 0.62.

For the multicolor experiment examining newly-eclosed adults 5d after heat-shock, the frequency of labeled ovarioles was 26/115 = 23%. If there were 20 target cells (likely in the range of the actual number of precursors), then p = 0.013 (because 0.987^20^= 0.77) and the frequency of single-cell lineages is 20×0.013x(0.987)^19^/0.23 = 0.86.

For the MARCM experiment examining newly-eclosed adults 5d after heat-shock, the frequency of labeled ovarioles was 33%. If there were 20 target cells, then p = 0.02 (because 0.98^20^= 0.67) and the frequency of single-cell lineages is 20×0.02x(0.98)^19^/0.33 = 0.80.

### Calculation of single-cell lineage frequencies assuming frequencies of single lineages based on the experimental frequency of ovarioles with marked ECs and FCs

In MARCM and multicolor lineage experiments scored in newly-eclosed adults we observed a higher frequency of ovarioles with labeled ECs and FCs but no FSCs (which we assume represent two distinct lineages because the marked cells are discontinuous) than would be expected based on the frequency of unmarked ovarioles and assuming all recombination events are independent. Based on the EC/FC clone frequency we used trial and error (using increments of 5%) to find the percentage of ovarioles with a single lineage that best fit the entire data-set of clone type frequencies we observed, using the simplifying assumption that all other marked ovarioles harbored exactly two lineages. In reality, the proportion of ovarioles with single lineages may be slightly different and a few ovarioles may include three or more marked lineages but the simplifying approximations we used are likely to be close to reality. These calculations, presented below, are better approximations than are generally made by simply considering the proportion of ovarioles with no marked cells.

#### MARCM clones induced 5d (0h APF) before eclosion

Single cell lineage frequencies and labeled cell content were calculated after estimating that 30% of marked ovarioles derived from a single marked cell, while the remaining 70% derived from two marked cells. In all cases, EC/FSC and FSC-only categories include clones with and without FCs (aggregated). pEC, pEC/FSC, pFSC, pFC are the proportions of each type of single-cell lineage amongst all marked lineages. Observed values are the proportion of marked ovarioles with the specific type(s) of cell. EC-only means no FSC. FC-only means no FSC. Both categories include EC+FC ovarioles.

Observed EC-only = 0.47 + 0.16 (with FCs) = 0.63 = 0.3 pEC + 0.7 (pEC)^2^ + 1.4pEC.pFC

Observed FC-only = 0.09 + 0.16 = 0.25 = 0.3 pFC + 0.7 (pFC)^2^ + 1.4pEC.pFC

Above is satisfied by pEC = 0.63 and pFC = 0.19

Observed FSC-only = 0.03 = 0.3pFSC + 0.7 [(pFSC)^2^ + 2pFSC.pFC] = 0.7[(pFSC)^2^ + 0.566pFSC So, pFSC = 0.050

Hence, pEC/FSC = 1 – (pEC + pFSC + pFC) = 0.13

The above values are the inferred single lineage frequencies of different types. The number of marked ECs and FSCs in each category is then deduced from these proportions and the observed number of marked cells in different types of marked ovariole.

Observed ECs per EC-only ovariole = (2.4 x0.47 + 2.9 x 0.16)/ 0.63 = 2.53

Of double-clones in EC-only, 0.397/0.636= 0.624 are deduced from clone type proportions to be two

EC clones and 0.239/0.636 = 0.376 are one EC and one FC clone

So, number of EC lineages per EC-only ovariole = 0.3 + 0.7(1.248 + 0.376) = 1.44

So ECs per EC-only lineage = 2.53/1.44 =1.76

Observed FSCs per FSC-only ovariole = 14/3

Of double-clones in FSC-only, 0.0025/0.215= 0.116 are two FSC clones and 0.019/0.215 = 0.884 are one FSC and one FC clone

So, number of FSC lineages per FSC-only ovariole = 0.3 + 0.7(0.232 + 0.884) = 1.08

So FSCs per FSC-only lineage = 14/(3×1.08) = 4.32

Total ovarioles with marked cells= 98, so total number of lineages= 98 x 1.7 = 166.6 Total ECs from EC-only lineages = 166.6pEC(1.76) = 184.7

Total FSCs from FSC-only lineages = 166.6pFSC(4.32) = 35.9

Total ECs scored = 253, total FSCs scored = 111 So, total ECs from EC/FSC clones = 68.3

Total FSCs from EC/FSC clones = 75.1

Total EC/FSC lineages = 166.6pEC/FSC = 21.66

So, ECs per EC/FSC clone = 3.15, and FSCs per EC/FSC clone = 3.47

Deduced 24.0 precursors (to produce 16 FSCs): 15.1 pECs, 3.1 pEC/FSCs, 1.2 pFSCs, 4.6 pFCs.

#### Multicolor clones induced 5d (0h APF) before eclosion

Analogous calculations for multicolor GFP-only clones where EC plus FC category was 8% and we consequently assume that 70% of marked ovarioles have lineages derived from one cell, and 30% have two lineages.

Observed EC-only = 0.423 + 0.077 = 0.50 = 0.7 pEC + 0.3 (pEC)^2^ + 0.6pEC.pFC

Observed FC-only = 0.077 + 0.077 = 0.154 = 0.7 pFC + 0.3 (pFC)^2^ + 0.6pEC.pFC

Above satisfied by pEC= 0.52 and pFC= 0.14

Observed FSC-only = 0.077 = 0.7pFSC + 0.3 [(pFSC)^2^ + 2pFSC.pFC] = 0.3[(pFSC)^2^ + 0.778pFSC

So, pFSC = 0.095

Hence, pEC/FSC = 1 – (pEC + pFSC + pFC) = 0.245

The above values are the inferred single lineage frequencies of different types. The number of marked ECs and FSCs in each category is then deduced from these proportions and the observed number of marked cells in different types of marked ovariole.

Observed ECs per EC-only ovariole = 38/13 = 2.92

Of double-clones in EC-only, 0.2704/0.416 = 0.65 are two EC clones and 0.1456/0.416 = 0.35 are one EC and one FC clone

So, number of EC lineages per EC-only ovariole = 0.7 + 0.3(1.3 + 0.35) = 1.195

So ECs per EC-only lineage = 2.92/1.195 = 2.44

Observed FSCs per FSC-only ovariole = 4.0

Of double-clones in FSC-only, 0.00593/0.02363= 0.25 are two FSC clones and 0.0177/0.02363= 0.75 are one FSC and one FC clone

So, number of lineages per FSC-only ovariole = 0.7 + 0.3(0.50 + 0.75) = 1.0375

So FSCs per FSC-only lineage = 4.0/1.0375 = 3.86

Total ovarioles with marked cells= 26, so total number of lineages= 26 x 1.3 = 33.8

Total ECs from EC-only lineages = 33.8pEC(2.44) = 42.89

Total FSCs from FSC-only lineages = 33.8pFSC(3.86) = 10.04

Total ECs scored = 66, total FSCs scored = 29

So, total ECs from EC/FSC clones = 23.11

Total FSCs from EC/FSC clones = 18.96

Total EC/FSC lineages = 33.8pEC/FSC = 8.28

So, ECs per EC/FSC clone = 2.79, and FSCs per EC/FSC clone = 2.27

Deduced 17.3 precursors (to produce 16 FSCs): 9.0pECs, 4.2 pEC/FSCs, 1.6 pFSCs, 2.0 pFCs.

#### MARCM clones induced 3.5d (36h APF) before eclosion

Analogous calculations for MARCM clones induced at −3.5d, where EC plus FC category was 44% and assuming all marked ovarioles have two lineages.

Observed EC-only = 0.207 + 0.438 = 0.645 = (pEC)^2^ + 2pEC.pFC

Observed FC-only = 0.091 + 0.438 = 0.529 = (pFC)^2^ + 2pEC.pFC

Above satisfied by pEC= 0.51 and pFC= 0.38

Observed FSC-only = 0.033 = (pFSC)^2^ + 2pFSC.pFC = (pFSC)^2^ + 0.76pFSC So, pFSC = 0.041

Hence, pEC/FSC = 1 – (pEC + pFSC + pFC) = 0.07

The above values are the inferred single lineage frequencies of different types. The number of marked ECs and FSCs in each category is then deduced from these proportions and the observed number of marked cells in different types of marked ovariole.

Of double-clones in FC-only, 0.144/0.532= 0.27 are two FC clones and 0.388/0.532 = 0.73 are one EC and one FC clone

Observed ECs per EC-only ovariole = 199/78 = 2.55

Of double-clones in EC-only, 0.260/0.646= 0.40 are two EC clones and 0.388/0.646 = 0.60 are one EC and one FC clone

So, number of EC lineages per EC-only ovariole = 0.8 + 0.6 = 1.40

So, ECs per EC-only lineage = 2.55/1.40 =1.82

Observed FSCs per FSC-only ovariole = 13/4 = 3.25

Of double-clones in FSC-only, 0.00168/0.03284= 0.05 are two FSC clones and 0.03116/0.03284= 0.95 are one FSC and one FC clone

So, number of FSC-only lineages per FSC-only ovariole = 1.05

So, FSCs per FSC-only lineage = 3.25/1.05 = 3.10

Total ovarioles with marked cells= 121, so total number of lineages= 242

Total ECs from EC-only lineages = 242pEC(1.82) = 224.6

Total FSCs from FC-only lineages = 242pFSC(3.10) = 30.8

Total ECs scored = 276, total FSCs scored = 111

So, total ECs from EC/FSC clones = 51.4

Total FSCs from EC/FSC clones = 80.2

Total EC/FSC lineages = 242pEC/FSC = 16.94

So ECs per EC/FSC clone = 3.03, and FSCs per EC/FSC clone = 4.73

Deduced 34.9 precursors (to produce 16 FSCs): 17.8pECs, 2.4 pEC/FSCs, 1.4 pFSCs, 13.3 pFCs. Also, by this time there are EC precursors that no longer divide (estimated at 5 for −4d and 18 for −3d from MARCM lineages examined in +2d adults, so perhaps 11.5 at −3.5d, to give a total of 45.4 precursors. So, estimated proportion of precursors that are pFCs is 13.3/45.4 = 29%.

#### bond-GAL4/G-trace lineages

Single lineage frequencies and content for clones initiated by *bond-GAL4* were calculated in a slightly different way (considering ovarioles with only marked ECs to calculate EC-only single lineage and then ovarioles with labeled ECs and FCs clones to calculate the FC-only lineage frequency) because there were almost no ovarioles with only marked FCs.

Here we assumed that every labeled ovariole had exactly two lineages.

Observed EC frequency = 38.36% = (pEC)^2^

So, pEC= 61.9%

Observed EC+FC frequency= 9.59% = 2pEC.pFC

So, pFC = 7.75%

Observed FSC frequency = 1.37% = (pFC)^2^ + 2pFC. pFSC

So, pFSC= 6.29%

Therefore pEC/FSC = 24.0%

Observed EC per EC-only (or EC+FC) lineage= 207/35= 5.91

Of double-clones in EC-only, expected frequencies: 0.383/0.479= 0.80 are two EC clones and 0.096/0.479 = 0.20 are one EC and one FC clone

So, number of EC lineages per EC-only ovariole = 2x 0.8 + 0.2= 1.8

So, ECs per EC-only lineage = 5.91/1.8 =3.28

Ovariole with labeled FSCs but no ECs has 7 FSCs

Of double-clones in FSC-only, 0.00396/0.01371= 0.289 are two FSC clones and 0.00975/0.01371 = 0.721 are one FSC and one FC clone

So, number of FSC lineages per FSC-only ovariole = 2×0.289 + 0.721 = 1.32

So, FSCs per FSC-only lineage = 7/1.32 = 5.3

Total ovarioles with marked cells= 73, so total number of lineages= 146

Total ECs from EC-only lineages = 146pEC(3.28) = 296.4

Total FSCs from FSC-only lineages = 146pFSC(5.3) = 48.7

Total ECs scored = 496, total FSCs scored = 190

So, total ECs from EC/FSC clones = 199.6

Total FSCs from EC/FSC clones = 141.3

Total EC/FSC lineages = 146pEC/FSC = 35.4

So, ECs per EC/FSC clone = 5.64, and FSCs per EC/FSC clone = 3.99

