## Supplemental Figures and Description of Movies for "Adult Stem Cells and Niche Cells segregate gradually from common precursors that build the adult Drosophila ovary during pupal development"

### Supplementary Materials

**Figure S1-S6 with legends.**

**Movies S1-S5.**

**Legends for Movies S1-S5:**

**Movie S1 to accompany Figure 9. A 40h APF multicolor germarium imaged for 11h 15min.** Clones were induced by heat shock such that cells could lose RFP, GFP or both, to yield green, red, or cells with no color, respectively. A green IC (white arrow) moved back and forth around the circumference of the germarium and stayed in its domain as the posterior cyst moved past. A red cell (red arrowhead) from a clone that began at the posterior of the posterior cyst moved along the cyst to its anterior. A green EGC cell (magenta arrow) appears in the z projections at 30 min and moved onto the posterior edge of the cyst. Two mitotic cells that divided at 2:30h (one in EGC and one in stalk) are indicated by white arrowheads.

**Movie S2 to accompany Figure 10. A 48h APF germarium imaged for 14h 45min.** 3D reconstruction of a movie of 48h APF *ubi-GFP*-labeled ovaries imaged every 15 min for 14h 45 min. The red, yellow, and pink cells described in Figure 10 and Figure S3C were tracked in Zeiss Zen software and then located in the 3D movie and colored using Procreate software.

**Movie S3 to accompany Figure 10. A 48h APF germarium imaged for 14h 45min.** The cell marked by the red asterisk in Figure 10 is shown here in a movie of individual z planes. The cell is tracked with a red arrow, and a white asterisk is present when this cell is undergoing mitosis. This cell started on the posterior of the posterior cyst and ended in the secondary EGC.

**Movie S4 to accompany Figure 10. A 48h APF germarium imaged for 14h 45min.** A movie of the individual z planes containing the cell marked by the white circle and asterisk in Figure 10 and Figure S4. This cell began in the basal stalk and ended on the budded cyst.

**Movie S5 to accompany Figure 11B. A 55h APF germarium imaged for 14h 45min.** A germarium at 55h APF with three tracked cells that began in the stalk and moved closer to the posterior cyst as cells from the EGC and stalk moved onto the cyst: the white cell began in the stalk 13 cells away from the cyst and ended in the stalk 4 cells from the cyst; the magenta cell began in the stalk 7 cells from the cyst and ended on the posterior of the cyst; and the yellow cell started as the most anterior cell in the stalk and ended on the posterior third of the cyst. At 3:00 the cell indicated by the blue arrow in the stalk went through mitosis.

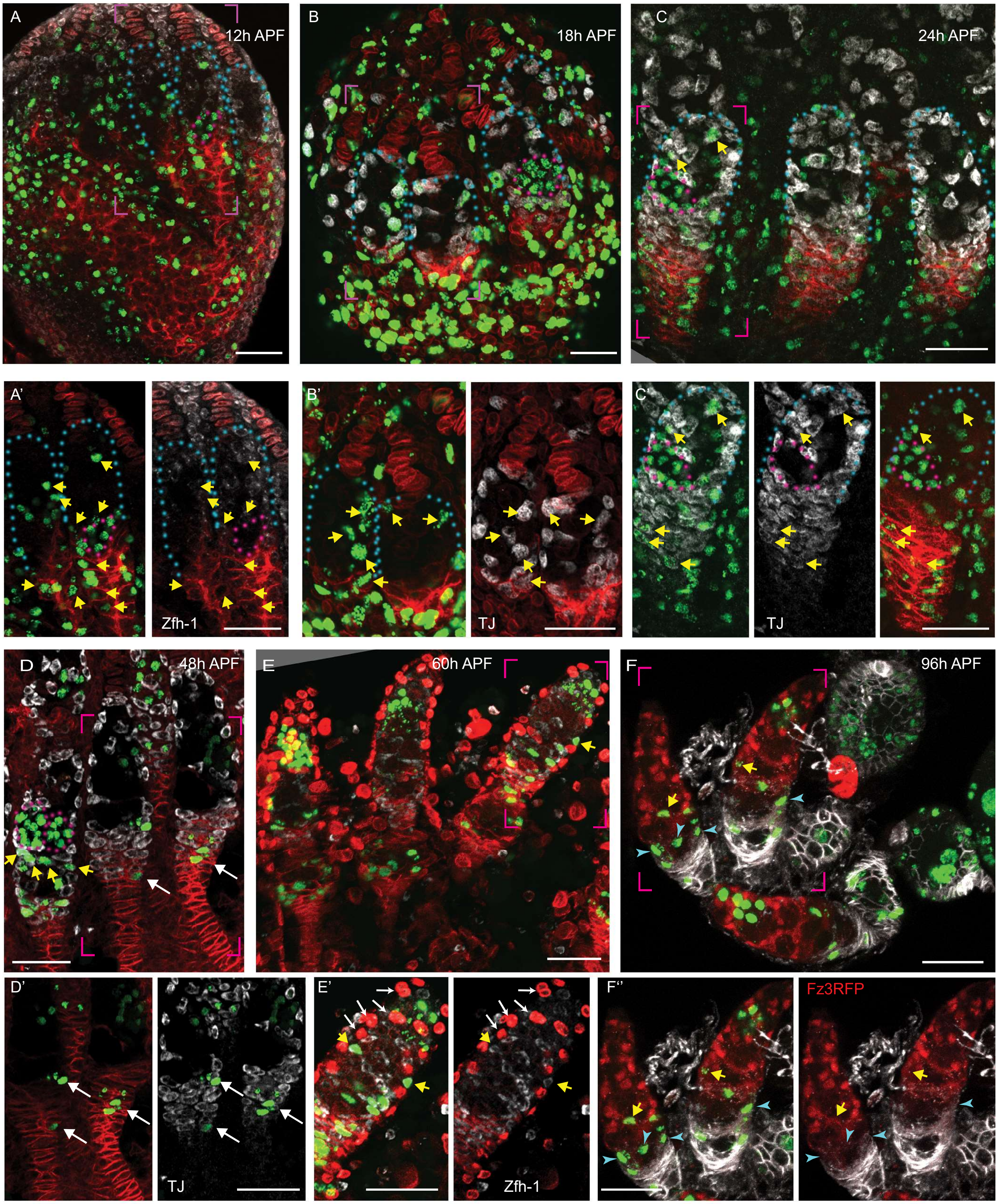


**Figure S1. Pattern of division of germline and somatic cells in pupal ovaries**

**Figure S1. Pattern of division of germline and somatic cells in pupal ovaries**

(A-F) Pupal ovaries from 0h to 96h APF were incubated with EdU (visualized in green) to label S-phase cells prior to fixation and staining for (A, E) Zfh-1 (white), (B-D) TJ (white) or (F) Fz3-RFP (red) to identify somatic cells, (A-E) LamC and Fas3 (both red), to mark TFs and basal stalks, respectively or (F) Fas3 only (white). Blue dotted lines outline individual germaria and pink bracketed regions are expanded in panels below each main image. All scale bars are 20 µm. (A-C) EdU (green) in the most posterior germline cyst (pink outlines) was seen at (A') 12h APF, (B) 18h APF and (C') 24h APF. (A-F) Somatic cells with (green) EdU (yellow arrows) were observed (A-C) among all ICs and at higher frequency in Fas3-positive cells posterior to the developing germarium from 12-24h APF. (D-F) ICs in the anterior third of the germarium were rarely labeled at 48h APF or later (green signals without yellow arrows are germline cells), but (D, D') TJ-positive, Fas3-positive EGC cells were frequently labeled (white arrows) and (E, E') a few ICs in the anterior half of the germarium (shown in different z-sections in E') were still labeled at 60h APF before (F) EdU labeling was confined to more posterior cells by 96h APF. (E, F) At 60h APF one egg chamber has budded and LamC stains epithelial sheath cells, visible also (F) at 96h APF after three egg chambers have budded. Here, many somatic cells are labeled with EdU close to a stage 2b germline cyst (blue arrowheads).

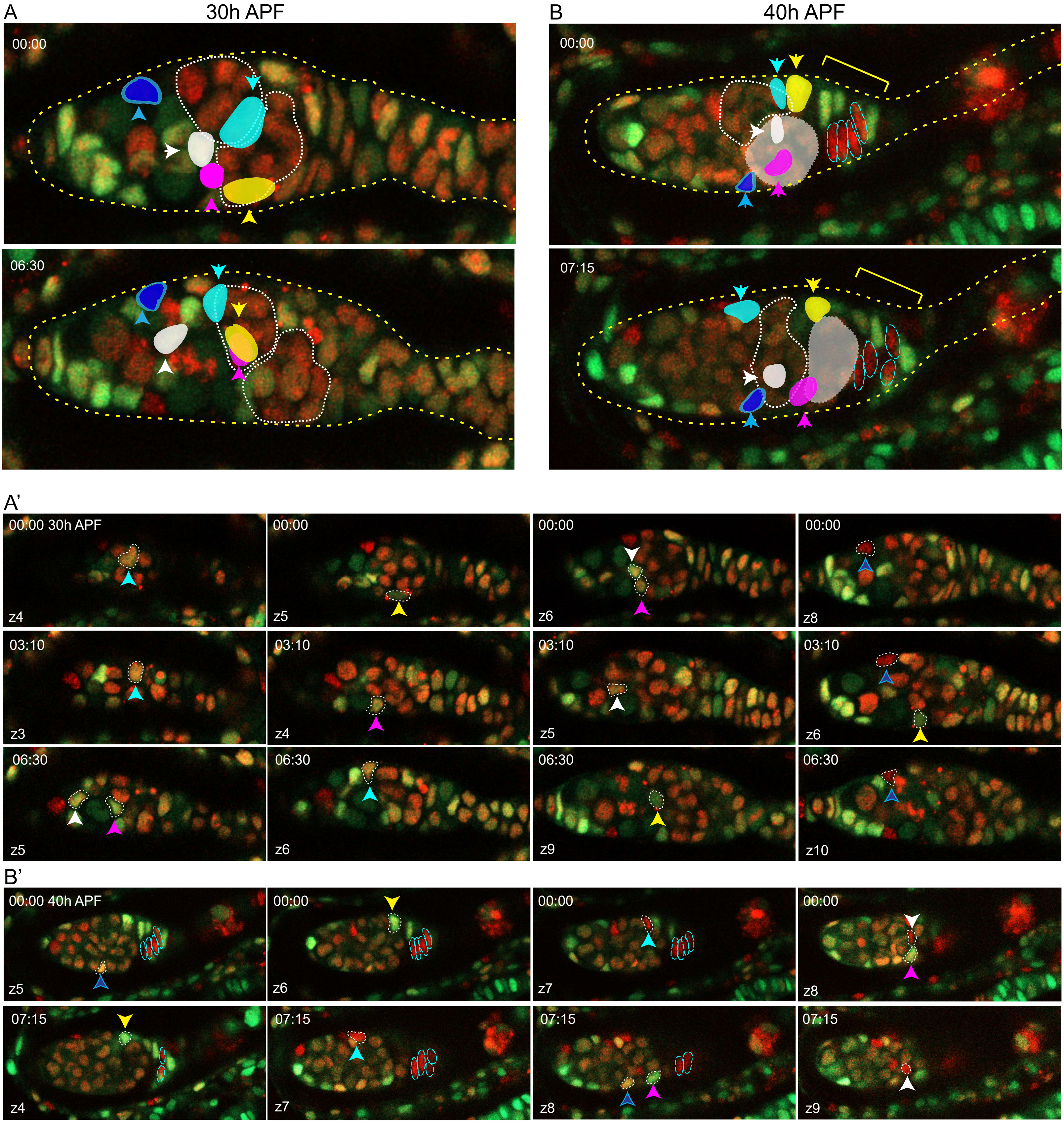


**Figure S2. ICs move independently of one another and do not mix with EGCs.**
(A) Z-projection of selected time-points from live imaging of a 30h-APF ovary for 6h 30min, showing five tracked ICs. ICs were located at the outer surface of the germarium and moved independently of one another in posterior, anterior, and lateral directions (around the circumference). The most posterior germline cyst (outlined by a white dotted line) moved posteriorly past the cyan and yellow IC cells. The relative disposition of ICs and cysts changed considerably, suggesting they are not strongly associated. Only z planes containing tracked cells were included in the projection (z4, 5, 6, 7, and 8 at 00:00 and z5, 6, 9, and 10 at 06:30) (A') Individual z sections of the 30h-APF ovary (arranged horizontally for each time point) show that ICs move around the circumference of the germarium. The cyan and yellow cells began on the bottom of the germarium and moved towards the mid-section, whereas the pink and white cells started in a mid-section and moved to the bottom.

(B) Z-projection of selected time-points from live imaging of a 40h-APF ovary for 7h 15min, showing five tracked ICs. ICs moved independently of one another and remained in their domain as the most posterior germline cyst (highlighted in white) moved past. Four cells in the EGC (outlined red cells) remained in the EGC (indicated by the yellow bracket). Projected sections are z5-9 at 00:00 and z4, 7, 8 and 9 at 07:15. (B') Individual z sections of the 40h-APF ovary show that IC cells move independently in all directions, and EGC cells did not move relative to one another, suggesting greater cohesion.

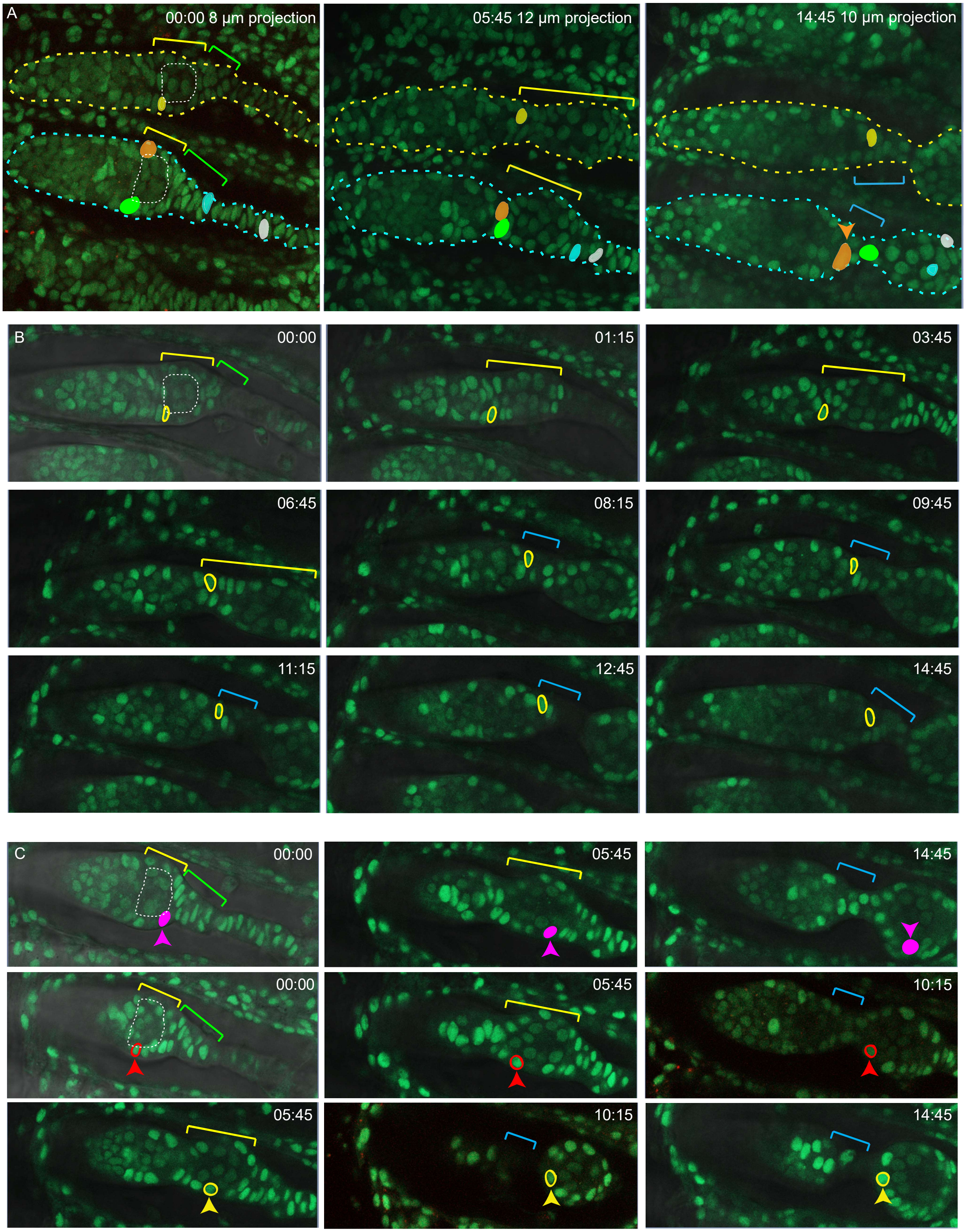


**Figure S3. Detail from live imaging (Movie S2) summarized in Figure 10.**

**Figure S3. Detail from live imaging (Movie S2) summarized in Figure 10.**

Selected time-points from a movie of a 48h APF ovary labeled with ubi-GFP, in which individual cells were tracked using Zeiss Zen through 60 time-points over 14h 45 min. Posterior cysts at the beginning of the movie are highlighted by a dotted white line. (A) Z projections from the beginning, middle and end of imaging. (B) Selected individual z slices from the movie show that the yellow cell in the upper germarium started just anterior to the posterior cyst and ended in the second EGC (magenta brackets). (C) Individual z slices from the lower germarium showing progress of the red, pink and yellow cells. The pink cell started close to the posterior face of the most mature germline cyst and ended near the middle of that budded cyst. The red cell started near the middle of the cyst and was near the anterior of the budded at cyst by 10h 15 min, after which it could not be tracked. The yellow cell was first tracked from 5h 45min when it was near the middle of the budded cyst. Nine hours later it was at the anterior edge of the budded cyst.

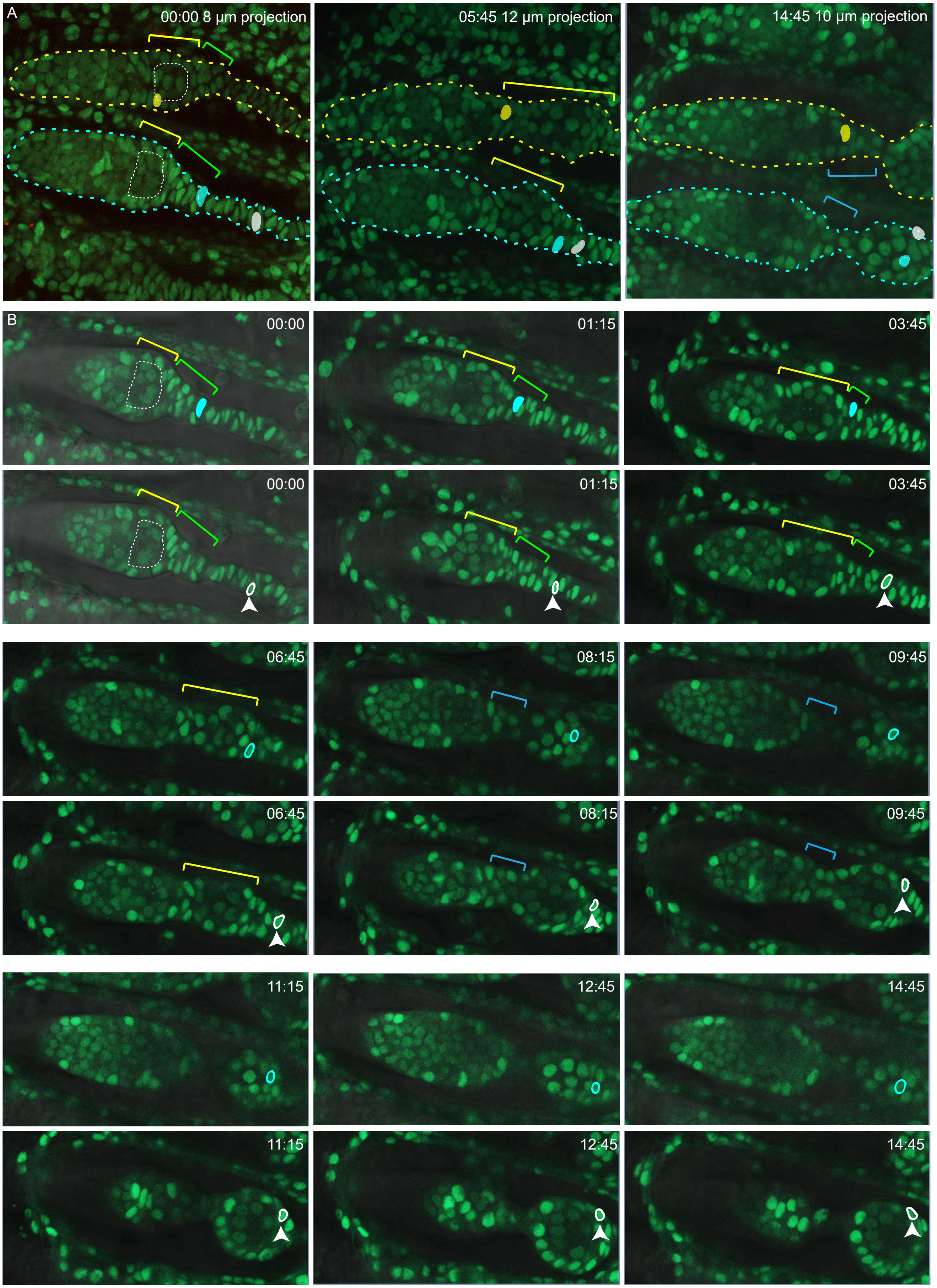


**Figure S4. Further detail from live imaging (Movie S2 and S4) summarized in Figure 10.**

**Figure S4. Further detail from live imaging (Movie S2 and S4) summarized in Figure 10.**

Time-points from a movie of a ubi-GFP-labeled 48h APF ovary in which individual cells were tracked using Zeiss Zen. (A) Z projections from the beginning, middle and end of imaging. (B) Individual z slices show that the cyan cell started in the EGC (yellow bracket) and ended on the posterior half of the budded cyst. The white cell started posterior to the EGC in basal stalk territory, became part of the EGC by 3h 45min and ended on the posterior half of the budded cyst.

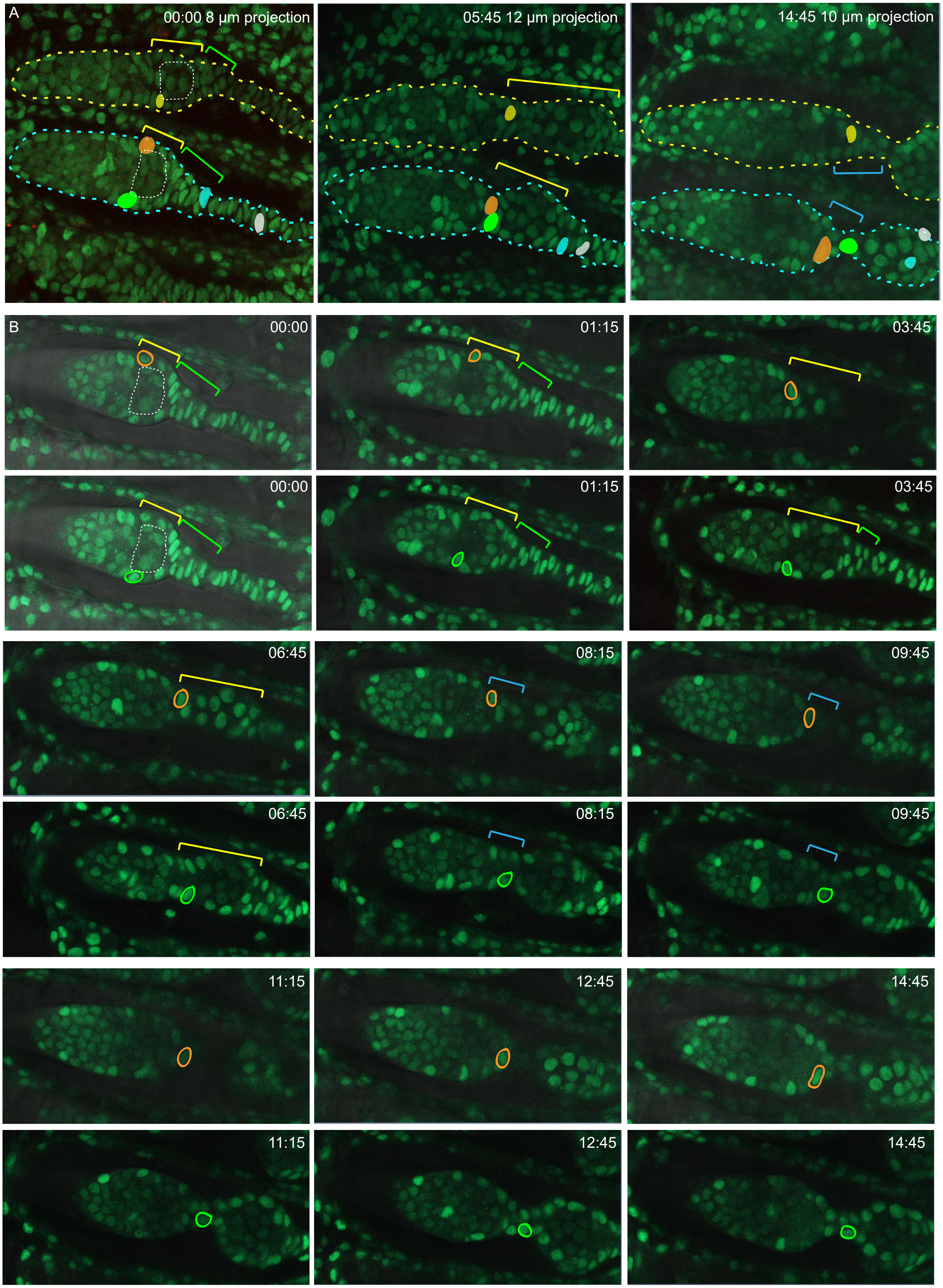


**Figure S5. Further detail from live imaging (Movie S2) summarized in Figure 10.**

**Figure S5. Further detail from live imaging (Movie S2) summarized in Figure 10.**

Time-points from a movie of a ubi-GFP-labeled 48h APF ovary in which selected cells were tracked using Zeiss Zen. (A) Z projections from the beginning, middle and end of imaging. (B) Individual z slices show that the green and orange cells started at the far anterior of the most posterior cyst. The green cell ended between the germarium and the budded cyst, while the orange cell ended at the posterior edge of the germarium.

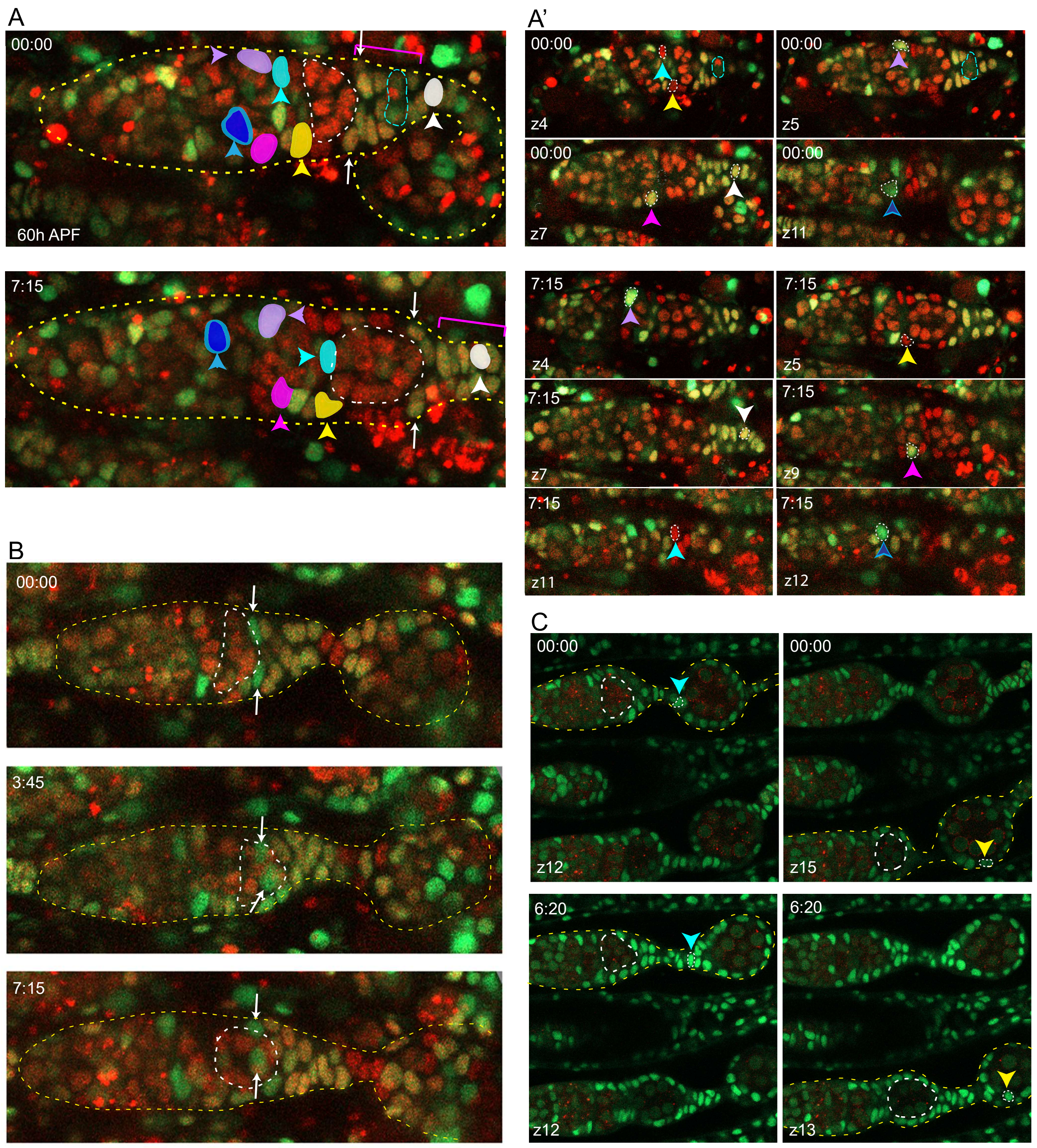


**Figure S6. Continued anterior movement of FCs after budding of the first egg chamber.**
Live imaging of 60h-APF germaria with one budded egg chamber, highlighted by yellow dotted lines, showed that FCs continue to move short distances anterior in relation to germline cysts. Germaria in A and B are from the same multicolor ovary. (A) Z projections of two timepoints show that tracked ICs (colored cells) remained anterior to the budding cyst as the germarium lengthened. Cells in the secondary EGC (indicated with a magenta bracket) moved a short distance anteriorly around the budding cyst (white arrows) and within the EGC (white colored cell) as the budding cyst moved posterior. Several cells in the secondary EGC and stalk divided. A mitotic cell is circled by the cyan dotted line at time 00:00. At 00:00, 4 z-planes (4, 5, 7, 11) are projected, and at 07:15, 6 z-planes (4, 5, 7, 9, 11,12) are projected. (A') Individual z slices show the movement of the cells along the z-axis. Time 00:00 is shown in the four z slices on top, with the cyan dotted line circling a dividing cell. ICs moved independently in all directions including around the circumference of the germarium. (B) A 60h APF multicolor germarium with two green cells (white arrows) that started posterior on the budding cyst and moved anterior along it as the cyst moved posteriorly. Many cells in the secondary EGC and stalk were dividing during this movie and the stalk can be seen to lengthen. (C) Cells were tracked in a movie of a 60h APF ovary labeled with ubi-GFP. Left panels: in the upper germarium a cell indicated by the cyan arrowhead moved anteriorly from a location on the budded cyst to the stalk, which can be seen to lengthen. Right panels: in the lower germarium, an FC indicated by the yellow arrowhead moved further anterior on the surface of the budded cyst.
